# Symbiosis between river and dry lands: phycobiont dynamics on river gravel bars

**DOI:** 10.1101/2020.01.29.924803

**Authors:** Lucie Vančurová, Veronika Kalníková, Ondřej Peksa, Zuzana Škvorová, Jiří Malíček, Patricia Moya, Kryštof Chytrý, Ivana Černajová, Pavel Škaloud

**Author notes:** Corresponding author: Lucie Vančurová, Charles University, Faculty of Science, Department of Botany, Benátská 2, 128 01 Prague 2, Czech Republic.

## Abstract

River gravel bars are dynamic and heterogeneous habitats standing on transition between aquatic and terrestrial environment. Periodical flooding, low nutrient content, frost, missing safe sites, drought, and heat on the ground surface significantly influence life in these habitats. Mutualistic symbiosis may be a successful strategy for organisms to survive and to proliferate under harsh conditions. The lichen genus *Stereocaulon* was selected as a model symbiotic system among the organisms living on river gravel bars. The aim of our work was to determine effect of this dynamic environment on a phycobiont (i.e., green eukaryotic photobiont) community structure. We analysed 147 *Stereocaulon* specimens collected in the Swiss Alps using Sanger sequencing (fungal ITS rDNA, algal ITS rDNA, algal actin type I gene) and 8 selected thalli and 12 soil samples using Illumina metabarcoding (ITS2 rDNA). We performed phytosociological sampling on each study plot (n=13). Our analyses of communities of phycobionts, lichens, bryophytes, and vascular plants indicated an ongoing colonisation by phycobionts and gradual change of phycobiont community along to the successional gradient. We recovered great phycobiont diversity associated with *Stereocaulon* mycobionts including algae reported as phycobionts for the first time. Each of two *Stereocaulon* mycobiont OTUs has distinct pool of predominant phycobionts in the study area. Finally, all thalli selected for Illumina metabarcoding contained a wide range of additional intrathalline algae, i.e., showed algal plurality. In general, succession process on newly emerged or recently disturbed localities also takes place within a community of microscopic symbiotic organisms, such as phycobionts.

## Introduction

River gravel bars are dynamic and heterogeneous habitats ranging from glacial floodplains and alpine wide river valleys to piedmont (Montgomery and Buffington 1998; Malard et al. 2006; Tockner et al. 2006; Hohensinner et al. 2018). Periodical floods together with variation of speed and intensity of the water current, model richly braided channels with a mosaic of channels, pools, bars and islands (Junk et al. 1989; Tockner et al. 2000; Ward et al. 2002). Destruction and reformation of the river gravel bars by floods results in a wide structural change and a cyclic vegetation succession. Gravel bar vegetation is characterized by early-successional pioneer vegetation and subsequent, mainly shrub, vegetation stages in the colonization sequence (Pettit and Froend 2001; Prach et al. 2016). These vegetation types form a mosaic of units according to microhabitat environmental conditions and disturbation level (Müller 1996; Tockner et al. 2000; Wellstein et al. 2003; Gilvear et al. 2008). Gravel bars in alpine zones contribute significantly to the regional diversity of an otherwise harsh alpine environment (Tockner and Malard 2003). Early-to mid-successional stands are often occupied by relatively species rich communities of vascular plants of high evenness and relatively low vegetation cover. In later successional stages, the evenness and species richness decrease as organic matter and nutrients accumulate and the competition from established dominant species increase. This pattern was shown gravel bars, for example, by Corenblit et al. (2009), Prach et al. (2014), Kalníková et al. (2018).

Alpine rivers with glacier source are highly influenced by daily flooding, due to melting of glaciers which makes the environment even more extreme (e.g. Milner and Petts 1994; Tockner et al. 2000; Malard et al. 2006). Low nutrient content in a substrate surface represents the most limiting environmental factor in the glacier floodplains. Especially the recently unglaciated terrain is characterized by bare soils which do not contain any organic matter and the soil seed bank is initially not present. In addition, frost, missing safe sites, drought, and heat on the ground surface significantly influence life in these habitats (Stöcklin and Bäumler 1996; Tockner et al. 2006; Marcante et al. 2014). Common strategies of the gravel bar species are usually high diasporas dispersability, fast growth, disturbance tolerance, clonal growth and ability to grow on poor soils (Jeník 1955; Vitt et al. 1986; Muotka and Virtanen 1995; Stöcklin 1999; Karrenberg et al. 2003; Ellenberg and Leuschner 2010).

Under harsh conditions of river gravel bars, mutualistic symbiosis may be a successful strategy for the organisms to survive and to proliferate (Doty et al. 2016). Since lichens represent one of the oldest known and recognizable examples of mutualistic symbiosis living under stressful conditions (Seckbach and Grube 2010), we selected the *Stereocaulon* lichens, among the organisms living on river gravel bars, as a model symbiotic system. The *Stereocaulon* genus is widespread and ecologically successful and is a pioneer lichen growing in harsh conditions on newly formed substrates (Stretch and Viles 2002; Meunier et al. 2014). Moreover, previous studies confirmed its ability to survive episodic submersion (Sadowsky et al. 2012), even though it is not natively aquatic. Recently, an exceptional diversity of phycobionts was discovered to be associated with *Stereocaulon*, including three ecologically diversified trebouxiophycean genera *Asterochloris, Vulcanochloris* and *Chloroidium* (Vančurová et al. 2015, 2018).

The lichen host ecological amplitude may be greatly influenced by its specificity towards the photobionts (Rolshausen et al. 2017; Vančurová et al. 2018). Symbiotic interactions vary along the environmental gradients (Godschalx et al. 2019) and could be affected by stressful environments (Romeike et al. 2002; Engelen et al. 2010). Therefore, the aim of our work was to determine the phycobiont diversity pattern of *Stereocaulon alpinum* in the gradient of the vegetation succession. We applied both Sanger sequencing (n=147) and Illumina metabarcoding (n=8) to *Stereocaulon alpinum* specimens collected in 13 study plots to address the following questions: 1) Is the phycobiont diversity influenced by succession, water regime, or alternatively other biotic and abiotic conditions?; 2) Do *Stereocaulon* lichens use their ability to co-operate with various phycobionts to cope with extreme conditions on river gravel bars of glacial floodplains?; 3) Do *Stereocaulon* lichens show algal plurality in this habitat?

## Material and Methods

### Study area and field sampling

The sampling was carried out in August 2017. The study area covered three glacial valleys (1) Morteratsch valley and 2) Roseg valley in the Bernina range and 3) Lötschental valley of the Lonza River in the Bernese Alps in the Swiss Alps. Four localities (Morteratsch, Roseg I, Roseg II and Lonza) situated on the river gravel bars of glacial floodplains (1995–2070 m a. s. l.) were sampled. We carried out 13 vegetation plots of 4 × 4 m. A detailed description of the location of each plot is provided in Online Resource 1/Table S1. Study plots in the localities belong to three succession stages: first (early-successional), second (defined by shrub layer), and third (the most developed usually with tree layer). In the locality Roseg II, only first and second stages were present. Examples of study plots belonging to different succession stages are given in Online Resource 2/Fig. S1.

Coordinates of each plot were recorded using a portable GPS device (WGS-84 coordination system). We recorded all lichen, bryophyte and vascular plant species within the vegetation plots. Lichens and bryophytes were collected from the ground and stones. In each plot we estimated visually: a) the cover of each species according to the extended Braun-Blanquet cover scale (Westhoff and Van Der Maarel 1978), b) the total vegetation cover and the cover of each layer (tree, shrub, herb, moss and lichen). We measured the elevation of the gravel bar (as a distance from its highest point to the actual water level with a tape) as well as distance from the river. One soil sample per plot was taken. Data on vegetation structure for each plot are listed in the Online Resource 1/Table S2. The substrate of all localities is rather acidic (in several cases with basic fractions), since the *Stereocaulon* lichens do not occur on calcareous substrata (Lamb 1951).

On each plot, at minimum 10 *Stereocaulon* samples were collected. For each *Stereocaulon* sample the type of substrate and affiliation to the plot was noted. Lichen morphospecies were identified in the field as well as in the laboratory using standard microscopic and chemical methods, including spot tests and thin-layer chromatography (TLC). *Stereocaulon* vouchers were deposited in Herbarium collection of the Charles University in Prague (PRC). Vouchers of accompanying lichens were deposited in the personal herbarium of J. Malíček. Vascular plants and bryophytes which were not identified in the field (unambiguous cases) were collected and identified in the lab. All records on vegetation plots were stored in the Gravel bar vegetation database – ID: EU-00-025 (Kalníková and Kudrnovsky 2017), which is included in the European Vegetation Archive (Chytrý et al. 2016). Nomenclature follows Euro+Med PlantBase (2006–2019) for vascular plants, Hill et al. (2006) for mosses, Grolle & Long (2000) for liverworts, and Nimis et al. (2018) for lichens.

### DNA extraction, amplification, and Sanger sequencing

DNA was extracted from lichen thalli (total lichen DNA). Lichen thalli were examined under a dissecting microscope and washed with water before DNA extraction to remove possible surface contamination. Total genomic DNA was isolated from thallus fragments following the CTAB protocol (Cubero et al. 1999). Both algal and fungal nuclear internal transcribed spacers (ITS rDNA) and the algal actin type I gene (including one complete exon and two introns located at codon positions 206 and 248; Weber and Kabsch 1994) were PCR amplified using primers listed in Online Resource 1/Table S3. PCRs were performed as described in Vančurová et al. (2018). All PCR were performed in 20 μl using Red Taq Polymerase (Sigma) as described by Peksa and Škaloud (2011) or with My Taq Polymerase. Negative controls, without DNA template, were included in every PCR run to eliminate false-positive results caused by contaminants in the reagents. The PCR products were sequenced using the same primers at Macrogen in Amsterdam, Netherlands. The newly obtained sequences were deposited in GenBank under accession numbers xxx (Online Resource 1/Table S4).

### Sequence alignment and DNA analyses

*Asterochloris* datasets were analyzed both as a single locus for the ITS rDNA (data not shown) and as a concatenated dataset of ITS rDNA and actin type I loci. The *Asterochloris* ITS rDNA dataset consisted of 202 sequences: 142 newly obtained sequences and 60 previously published sequences from *Stereocaulon* and other lichens retrieved from GenBank. The actin type I dataset consisted of 68 sequences: 7 newly obtained sequences, and 60 previously published sequences. The alignment was automatically performed by MAFFT v.7 software (Katoh and Standley 2013) under the Q-INS-I strategy and manually edited according to the published secondary structures of ITS2 rDNA (Škaloud and Peksa 2010) using MEGA v.6 (Tamura et al. 2013). The actin type I sequences were aligned using MAFFT v.7 software (Katoh and Standley 2013) under the Q-INS-I strategy. After deleting identical sequences, the resulting concatenated alignment comprised 64 samples represented by unique ITS rDNA and actin type I sequences.

The *Stereocaulon* (mycobiont) ITS rDNA dataset comprised 171 sequences: 145 newly obtained sequences and 26 representative sequences selected to cover all main clades 1–8 published by Högnabba (2006). The alignment was automatically performed by MAFFT v.7 software (Katoh and Standley 2013) under the Q-INS-I strategy. After removing identical sequences, the resulting alignment comprised 48 sequences. All DNA alignments are freely available on Mendeley Data: http://dx.doi.org/10.17632/jchg5h3t5k.1.

Phylogenetic relationships were inferred with Bayesian Inference (BI) carried out in MrBayes v.3.2.2 (Huelsenbeck and Ronquist 2001), maximum likelihood (ML) analysis implemented in GARLI v.2.0 (Zwickl 2006), and maximum parsimony (MP) analysis using PAUP v.4.0b10 (Swofford 2003). BI and ML analyses were carried out on a partitioned dataset to differentiate among ITS1, 5.8 S and ITS2 rDNA, actin intron 206, actin intron 248, and actin exon regions. The best-fit substitution models (Online Resource 1/Table S5) were selected using the Bayesian information criterion (BIC) implemented in JModelTest2 (Guindon and Gascuel 2003; Darriba et al. 2012). ML analysis was carried out using default settings, five search replicates, and the automatic termination set at 5 million generations. The MP analysis was performed using heuristic searches with 1000 random sequence addition replicates and random addition of sequences (the number was limited to 10^4^ per replicate). ML and MP bootstrap support values were obtained from 100 and 1000 bootstrap replicates, respectively. Only one search replicate was applied for ML bootstrapping.

### Statistical analyses

From the total of 147 samples with successfully sequenced phycobiont, two were excluded because of the absence of the mycobiont sequence. The phylogeny of *Stereocaulon alpinum* showed two lineages (OTU35 and OTU2; see below). Since the mycobiont identity affects the phycobiont diversity (Vančurová et al. 2018) four samples belonging to the minority species-level lineage (OTU2) were also excluded. Thus, statistical analyses were carried out using 141 members of the prevailing mycobiont species-level lineage (OTU35) and their photobiots.

The relationship between species richness of *Stereocaulon alpinum* (OTU35) phycobionts and overall lichen species richness was inspected as a correlation between the number of phycobiont species-level lineages and number of lichen species. Since the number of samples per plot varied, the number of phycobiont species-level lineages were down-sampled to the smallest sample size in the data set which was 5 samples (Online Resource 2/Fig. S2). After excluding the study plot number 11 (with sample size of 5 samples), the smallest sample size in the data set increased to 10 samples. The rarefaction was performed using the *rarefy* function in *vegan* R package (Oksanen et al. 2019). The linear regression was performed separately for dataset including all plots and for dataset restricted to plots with sample size ≥ 10. Since the parametric regression analyses can be significantly biased in small sample sizes, we performed the Bayesian linear regression instead. We constructed a regression model where we modelled the number of phycobionts (*X*_*i*_) as *X*_*i*_ ∼ *Normal* (*μ*_*i*_,*σ*), where *<u>μ</u>*_*<u>i</u>*_ was determined as *a + b * number of lichen species*_*i*_ (*a* = intercept, *b* = slope of the regression line) and *σ* as the variance of the residuals. The priors were set as follows: *a* ∼ Normal (0, 0.001), *b* ∼ Normal (0, 0.001), *σ* ∼ Uniform (0, 100).

The gradual change of phycobiont community composition was inspected as a correlation between the proportion of the two most abundant phycobiont species-level lineages ((number of *Asterochloris phycobiontica* samples + number of StA5 lineage samples)/number of all samples) and the succession stage (coded 1, 2, 3). We constructed a regression model where we modelled the proportion of the most abundant phycobionts (*X*_*i*_) as *X*_*i*_ ∼ *Normal* (*μ*_*i*_,*σ*), where *<u>μ</u>*_<u>*i*</u>_ was determined as *a + b * succession stage*_*i*_ (*a* = intercept, *b* = slope of the regression line) and *σ* as the variance of the residuals. The priors were set as described above. We ran three chains of the model for 1,000,000 iterations, discarding the initial 100,000 as burnin. We fit the regression model in program JAGS v. 4.2.0 (Plummer 2003) through the *R2JAGS* package (Su and Yajima 2009) in R.

To inspect the dynamics of phycobiont communities in the context of the river gravel bar habitats, two PCoA ordinations were created. We used two different proxies to explain the vegetation succession on river gravel bars: the cover of shrub layer and the cover of crustose lichens. For this purpose, successional status of study plots was classified into two groups (early successional and developed). This classification provides two alternative views on successional processes. Both ordination analyses were based on the square-rooted Bray-Curtis dissimilarity. The original Braun-Blanquet cover-abundance values were transformed to percentage and square-root transformed in order to calculate the dissimilarity index. The first ordination was computed based on vascular plant communities and the second on bryophytes + lichens. In both ordination plots, Ellenberg-type indicator values (Ellenberg et al. 1991) for light and temperature were passively superimposed. In the plot based on bryophytes + lichens, indicator values for bryophytes were used (Hill et al. 2006). We also compared if the variation of different taxa groups corresponds to similar environmental conditions, by calculating procrustes distances between taxa groups. The output was then used as input distance matrix for PCoA ordination, visualizing a relative dissimilarities of community composition gradients of the taxa groups.

The vegetation plot data were stored in Turboveg for Windows v.2 database (Hennekens and Schaminée 2001) and further managed with JUICE software (Tichý 2002) and in the R environment (R Core Team 2017) with the help of the *vegan* R package (Oksanen et al. 2019).

Bipartite association network between study plots and phycobiont species-level lineages was produced using *bipartite* R package (Dormann et al. 2008).

### Illumina metabarcoding of algal communities in selected lichen thalli and soil samples

In order to describe algal plurality in *Stereocaulon* thalli, Illumina metabarcoding was performed. Eight selected thalli (four assigned to mycobiont OTU35 and four to OTU2) were rehydrated with Milli-Q sterile water one day before being processed and stored in a growth chamber at 20°C under a 12h/12h light/dark cycle (15 µmol/m^2^/s). Thalli were cleaned under a stereomicroscope to remove soil particles and then superficially sterilized following Arnold et al. (2009). Fragments from different parts of each thallus were randomly excised and pooled together (0.1 mg). Total genomic DNA was isolated and purified using the DNeasy Plant Mini Kit (Qiagen, Hilden, Germany).

Soil from study plots (one sample per plot) was sampled. Soil samples were sieved to remove all possible contaminations. Total genomic DNA was isolated and purified using the Soil DNA Isolation Plus Kit^®^ (Norgen Biotek Corp.), following the manufacturer’s instructions.

Chlorophyta algal communities associated with the eight thalli and twelve soil samples were assayed using Illumina high-throughput sequencing of ITS2 of the rRNA operon, proposed as a universal barcode across eukaryotic kingdoms (Coleman 2009). High-coverage PCR primers at conserved sites were designed using a custom database for the algal phylum Chlorophyta (Online Resource 1/Table S3).

Amplicons for Illumina MiSeq sequencing were generated from nested PCR: in the first PCR the forward 1378-Chlorophyta (newly designed; Online Resource 1/Table S3) and the reverse ITS4 primers (White et al. 1990) were used (27 amplification cycles were run), in the second PCR three replicates were ampyfied using the primers 5.8F-Chlorophyta (newly designed; Online Resource 1/Table S3) and ITS4 modified with Illumina overhang adaptors (forward overhang: 5’-TCG TCG GCA GCG TCA GAT GTG TAT AAG AGA CAG-3’; reverse overhang: 5’-GTC TCG TGG GCT CGG AGA TGT GTA TAA GAG ACA G-3’; 22 cycles). These three replicates were then pooled together. PCR reactions were performed as described in Moya et al. (2017).

PCR products were purified using AMPure XP beads (Beckman Coulter). Indexing PCR and addition of Nextera sequence adapters were perfomed using Nextera XT Index kit (Illumina Inc., San Diego, CA, United States) following the protocol for Illumina L library preparation. Finally, a second purification round was carried out using AMPure XP beads. Libraries were then quantified and pooled together. The libraries were sequenced on Illumina MiSeq platform using the MiSeq Reagent Kit v3 (paired end 2× 300 bp), at STAB Vida, Lisbon, Portugal and Genomics Core Facility at the University of Valencia, Spain.

#### Bioinformatic analyses

Quality control analysis of the Illumina MiSeq paired-end reads was performed using the FastQC v.0.11.8. Raw reads were processed using Quantitative Insights Into Microbial Ecology 2 (QIIME2 v.2018.11; Bolyen et al. 2018). Demultiplexed paired-end sequence reads were preprocessed using DADA2 (Callahan et al. 2016), a package integrated into Qiime2 that accounts for quality filtering, denoising, joining paired ends, and removal of chimeric sequences. The first 20 bp were trimmed from forward and reverse reads before merging to remove adaptors. In order to remove lower quality bases, amplicon sequence variants (ASVs) were truncated at position 210 based on the FastQC reports during this step.

Subsequent analyses were based on the ASV table, which contained the count for each unique sequence in each sample. Only ASVs with frequency ≥100 were further analyzed. BLAST searches were used to confirm the sequence identity. Solely algal sequences were further analyzed. Phylogenetic tree (Online Resource 2/Fig. S3) was inferred with Bayesian Inference (BI) using MrBayes v.3.2.2 (Huelsenbeck and Ronquist 2001) as described above. Euler diagrams were produced using *eulerr* R package (Larsson 2019).

## Results

### Vegetation on study plots

In total, we noted 88 taxa of vascular plants, 19 taxa of bryophytes and 45 taxa of lichens within 13 study plots. Online Resource 1/Table S1 contains summary of species richness for particular plots.

### Phycobiont and mycobiont phylogenetic analysis

The predominant phycobiont detected in *Stereocaulon* thalli by Sanger sequencing of ITS rDNA, belong in a 97% of samples to the genus *Asterochloris* and only five specimens represent other trebouxiophycean algae. In the case of these five specimens, coded as A574, A574.1, A633, A634 and A634V the identity of the phycobiont was confirmed by Blast search against the GenBank database. Significant matches of 99% to 87% were obtained with *Coccomyxa viridis* HG973000 for A574, *Elliptochloris reniformis* LT560354 for A574.1 and uncultured Trebouxiophyceae FJ554399 for A633, A634 and A634V. These three sequences formed a well-supported clade with more distantly related sequence KF907701 (86% sequence similarity; Online Resource 2/Fig. S3) previously assigned to clade URa28 (Ruprecht et al. 2014). We follow this nomenclature hereafter for these sequences.

The phylogenetic hypothesis resulting from Bayesian analysis of the ITS rDNA and actin type I sequences of *Asterochloris* (Fig. 1) is congruent with previous studies (Peksa and Škaloud 2011; Gauslaa et al. 2013; Škaloud et al. 2015; Moya et al. 2015; Vančurová et al. 2018). The species boundaries delimited in Vančurová et al. 2018 and nomenclature used *ibidem* were maintained. We recovered a total of 14 lineages, including one novel lineage here referred to as StA9. Eleven of these lineages were previously found as phycobionts of *Stereocaulon* (Vančurová et al. 2018 and references therein), namely, *Asterochloris glomerata, A. irregularis, A. italiana, A. lobophora, A. phycobiontica, A.* aff. *italiana, Asterochloris* clade 8, *Asterochloris* clade 12, *Asterochloris* StA3, *Asterochloris* StA4 and *Asterochloris* StA5. Two (*Asterochloris echinata* and *A. leprarii*) were found in association with *Stereocaulon* mycobiont for the first time here. The most frequently occurring phycobionts linked with the lineages *Asterochloris* StA5 and *A. phycobiontica*.

**Fig. 1.**
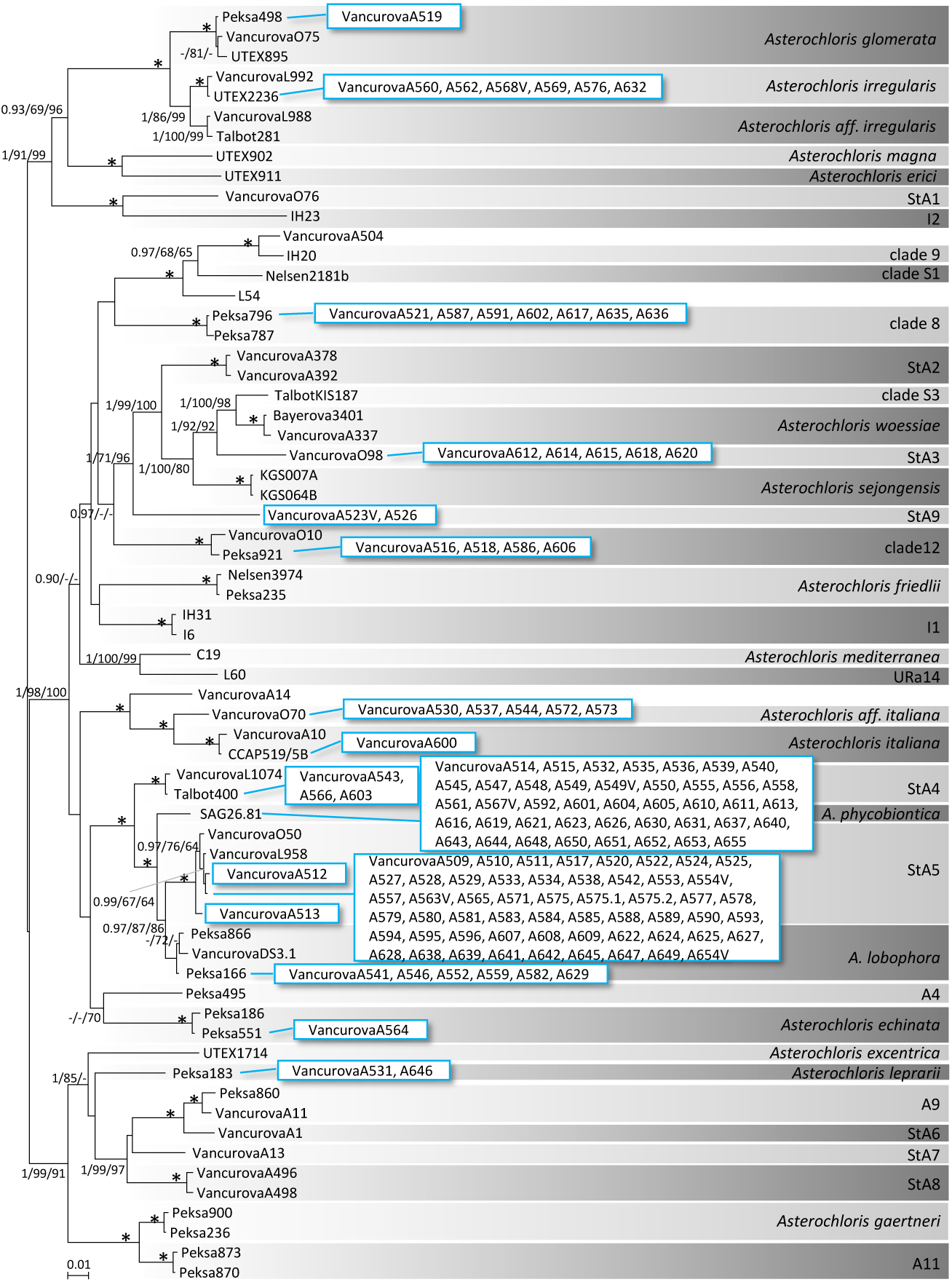
Phylogenetic hypothesis of *Asterochloris* resulting from the Bayesian analysis of combined ITS rDNA and actin type I sequences. Values at the nodes indicate the statistical supports of Bayesian posterior probability (left), maximum-likelihood bootstrap (middle) and maximum parsimony bootstrap (right). Fully supported branches (1.0/100/100) are marked with an asterisk. Scale bar shows the estimated number of substitutions per site. Newly obtained sequences are in boxes. Clade affiliations: clade 8, clade 9 *sensu* Škaloud and Peksa (2010), A4, A9, A11 *sensu* Peksa and Škaloud (2011), URa14 *sensu* Ruprecht et al. (2014), I1, I2 *sensu* Řídká et al. (2014), S1, S3 *sensu* Nelsen and Gargas (2006), *A.* aff. *irregularis, A.* aff. *italiana* and StA1 – StA8 *sensu* Vančurová et al. (2018). StA9 lineage was identified as new in present study. Online Resource 1/Table S6 contains accession numbers of reference sequences retrieved from GenBank

A phylogram resulting from Bayesian analysis of ITS rDNA sequences of *Stereocaulon* mycobiont is shown in Fig. 2. The majority of the recovered mycobiont sequences formed a well-supported lineage delimited as OTU35 by Vančurová et al. (2018). Five sequences matched with a distant related OTU2 (sister to DQ396973 and DQ396974), despite the morphological similarity of all studied samples. Both OTU35 and OTU2 fall into Group 8b *sensu* Högnabba (2006).

**Fig. 2.**
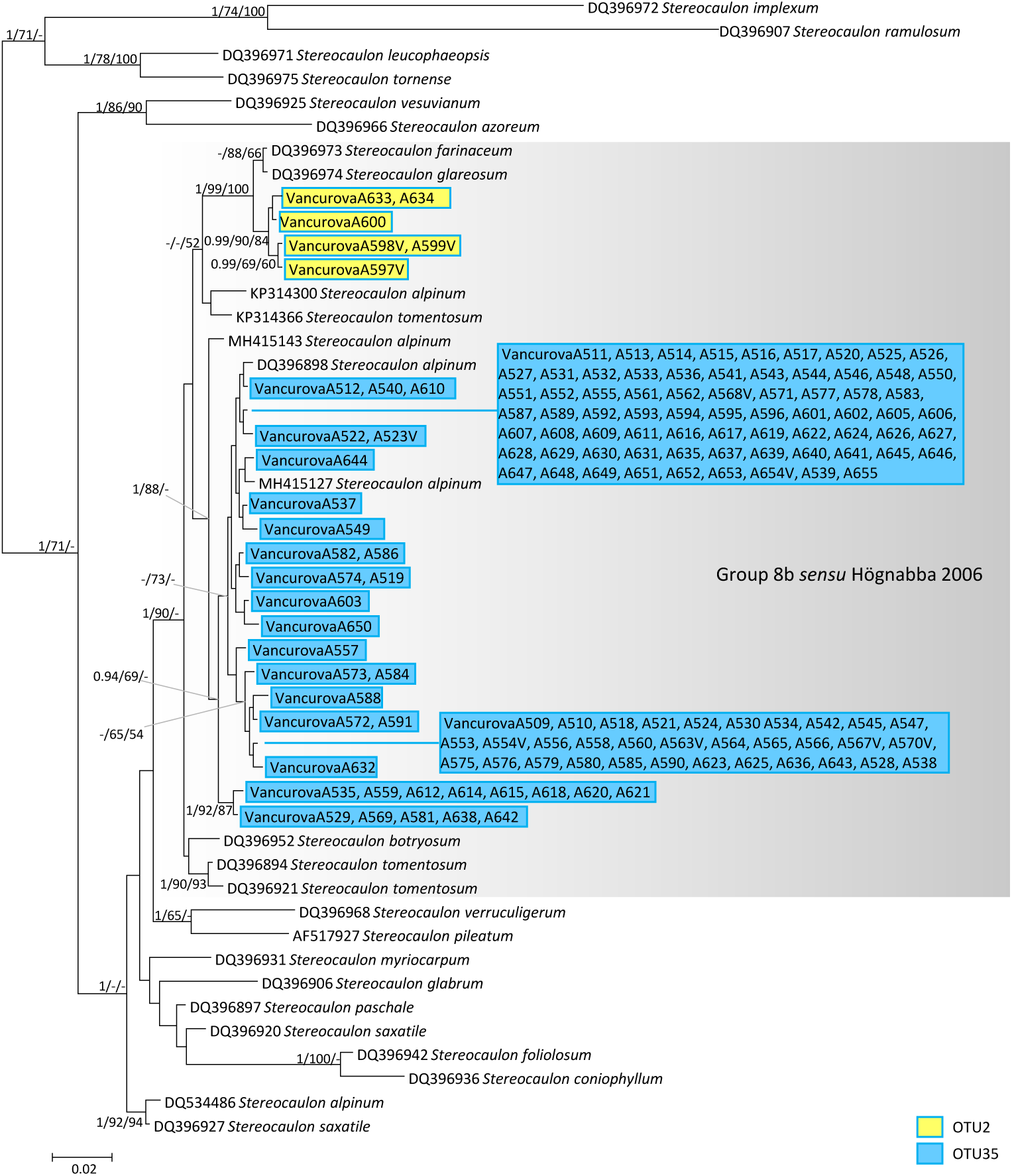
Phylogenetic hypothesis of *Stereocaulon* resulting from the Bayesian analysis of ITS rDNA. Values at the nodes indicate the statistical supports of Bayesian posterior probability (left), maximum-likelihood bootstrap (middle) and maximum parsimony bootstrap (right). Scale bar shows the estimated number of substitutions per site. Newly obtained sequences are in boxes. All new sequences are belonging to Group 8b *sensu* Högnabba (2006), as marked

### Phycobiont community structure and its changes

*Stereocaulon alpinum* OTU2 was sampled only in two study plots (numbers 8 and 11) and its phycobionts were successfully sequenced in four cases. One sequence was assigned to *Asterochloris italiana* and three sequences to trebouxiophycean lineage termed URa28. None of them was shared with the predominant *Stereocaulon* mycobiont (OTU35) in the study area.

*Stereocaulon alpinum* OTU35 (n=141) associated with 15 species-level lineages of phycobionts in the study area: 13 lineages of *Asterochloris*, one *Coccomyxa* and one *Elliptochloris*. Phycobionts from two to six species level lineages were recorded in each of 13 study plots (Fig. 3). When the sample size was reduced to five samples, 2-3.7 phycobiont species per plot were estimated. For the sample size of 10 (plot 11 with only five samples was excluded), 2.8-5.3 phycobiont species per plot were estimated.

**Fig. 3.**
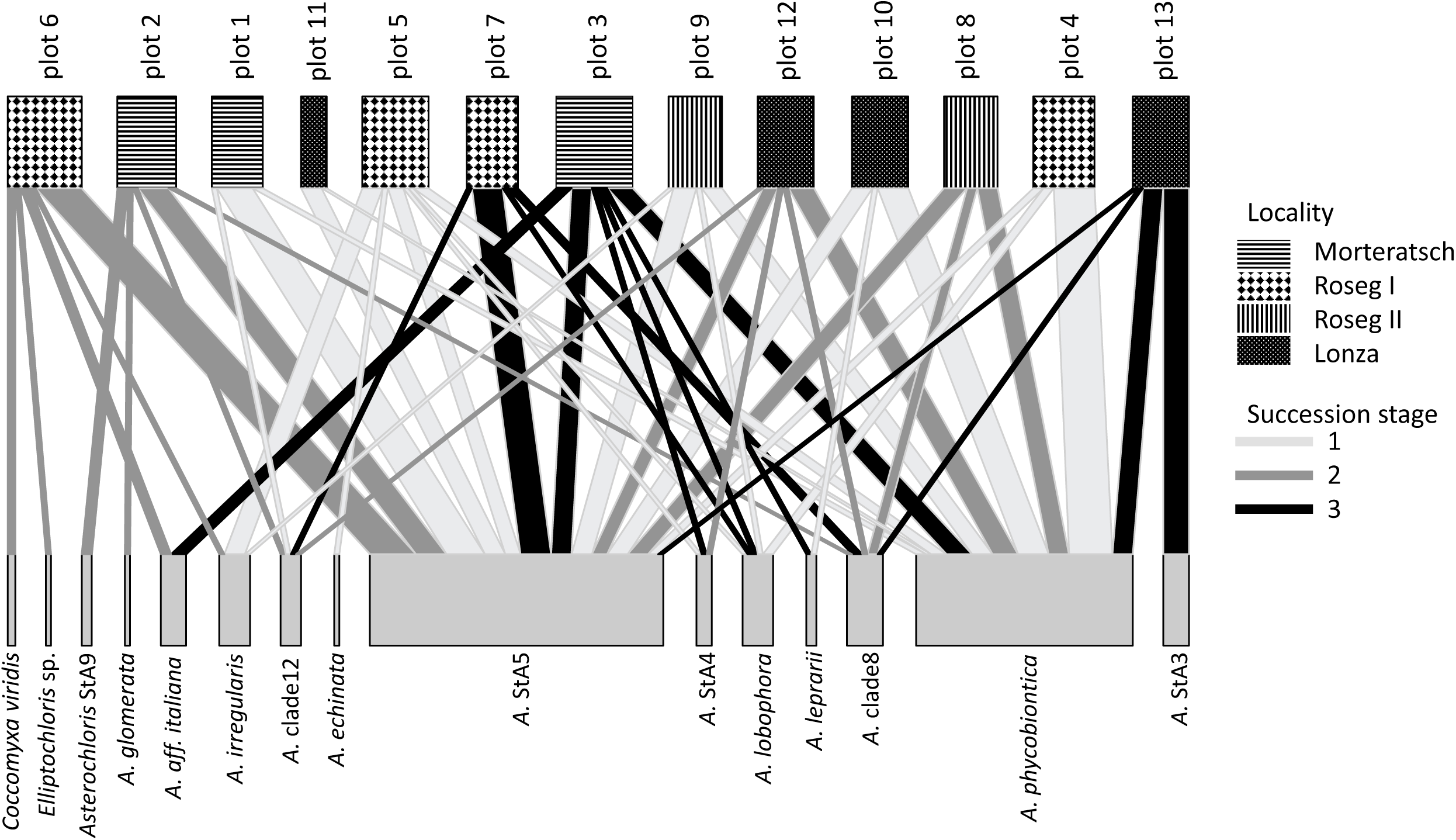
Bipartite association network between study plots and phycobiont species-level lineages. The width of the links is proportional to the number of specimens forming the association

Species richness of *Stereocaulon alpinum* OTU35 phycobionts increased significantly with the species richness of all lichens recorded at a study plot (Fig. 4). Moreover, gradual change of phycobiont community composition was proved using Bayesian linear regression (Fig. 5). The proportion of two most abundant phycobiont species-level lineages (*Asterochloris phycobiontica* and *Asterochloris* StA5) decreased significantly with the succession stage in favor of other species recovered in lower frequency.

**Fig. 4.**
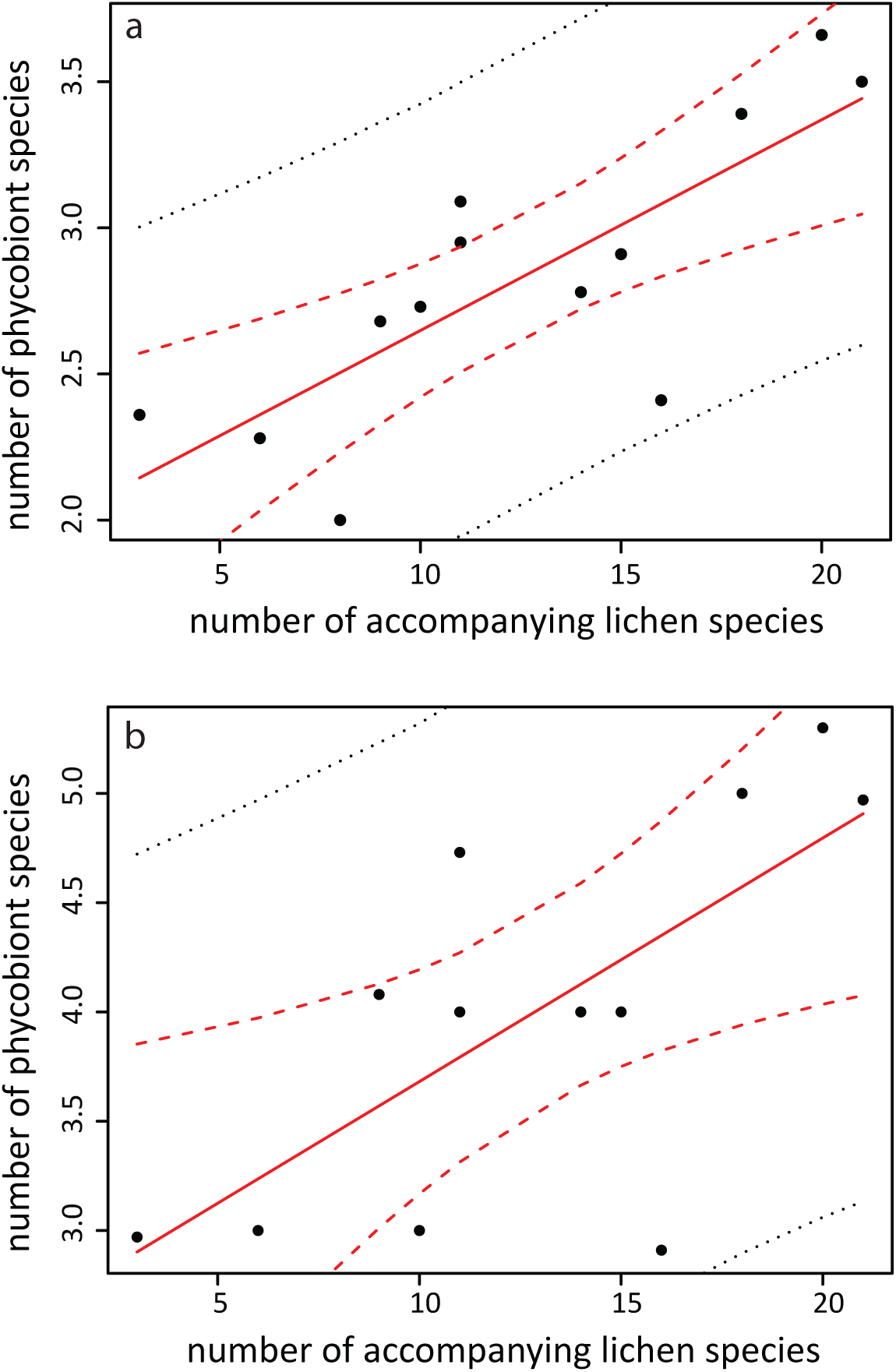
Bayesian linear regression of number of accompanying lichen species as a predictor of the number of phycobionts associated with mycobiont OTU35 down-sampled to **a** sample size of 5, **b** sample size of 10 per plot. Dashed lines show the 95% CRI around the regression line

**Fig. 5.**
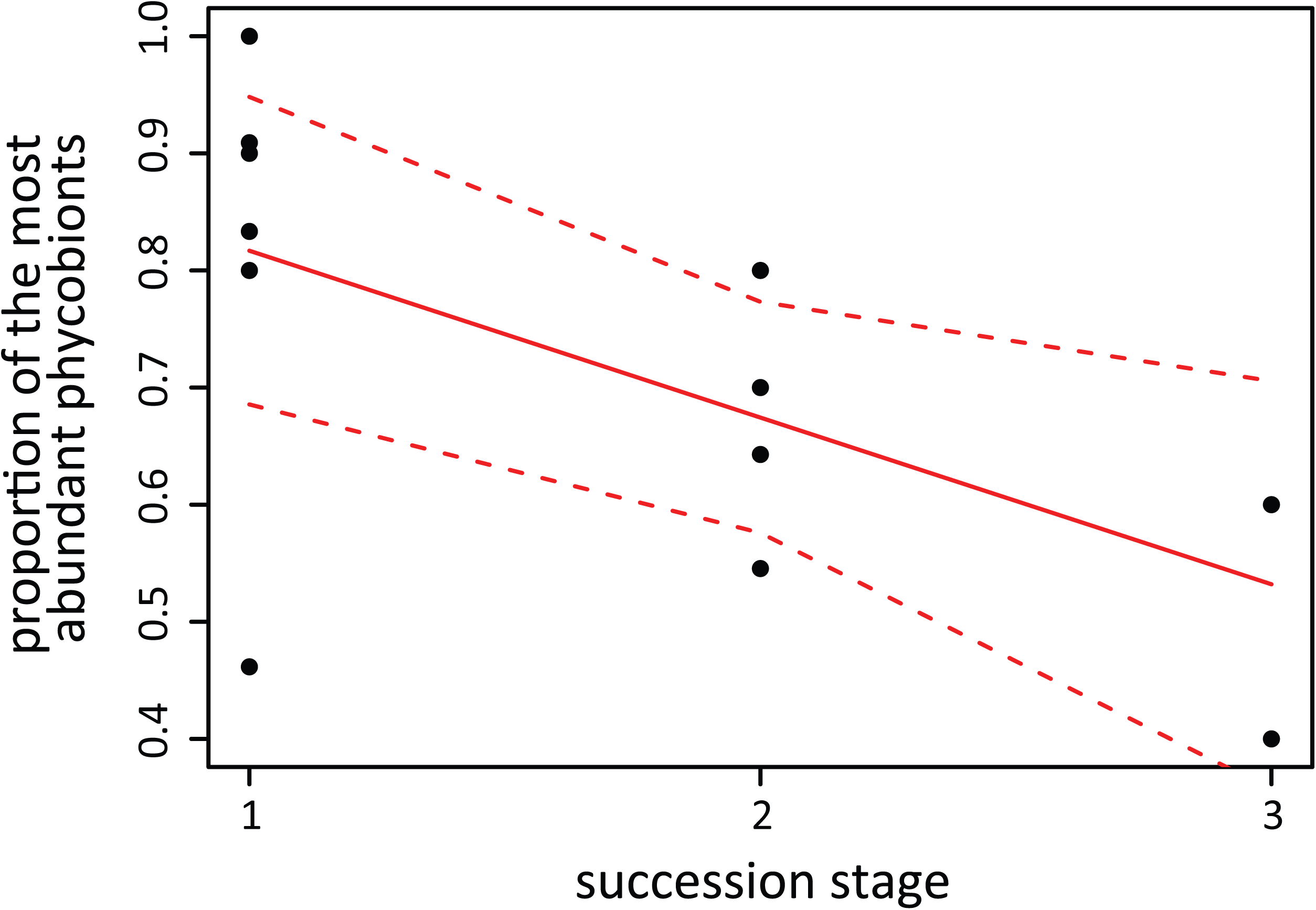
Bayesian linear regression of number of a succession stage as a predictor of the proportion of the most abundant phycobiont species-level lineages ((number of *Asterochloris phycobiontica* samples + number of StA5 lineage samples)/number of all samples). Dashed lines show the 95% CRI around the regression line

The gradient of succession in both ordinations (vascular-plants-based and non-vascular-plant-based; Fig. 6) positively correlated with the diversity of lichens and phycobionts and the number of locally rare phycobionts (defined as all species with the exception of the species *Asterochloris* StA5 and *A. phycobiontica* highly prevalent in the study area; Fig. 3). Phycobiont communities of early-successional stages were relatively species poor and mostly consisted of species generally abundant in the study area (thus, the species composition was similar within study sites). With ongoing succession, the number of locally rare phycobiont species increased altogether with the total number of phycobiont species and the number of lichen species. The diversity of vascular plants and bryophytes was not associated with this gradient. Higher moss layer cover and higher amount of light demanding species of both bryophytes and vascular plants were typical of the early-successional stages.

**Fig 6.**
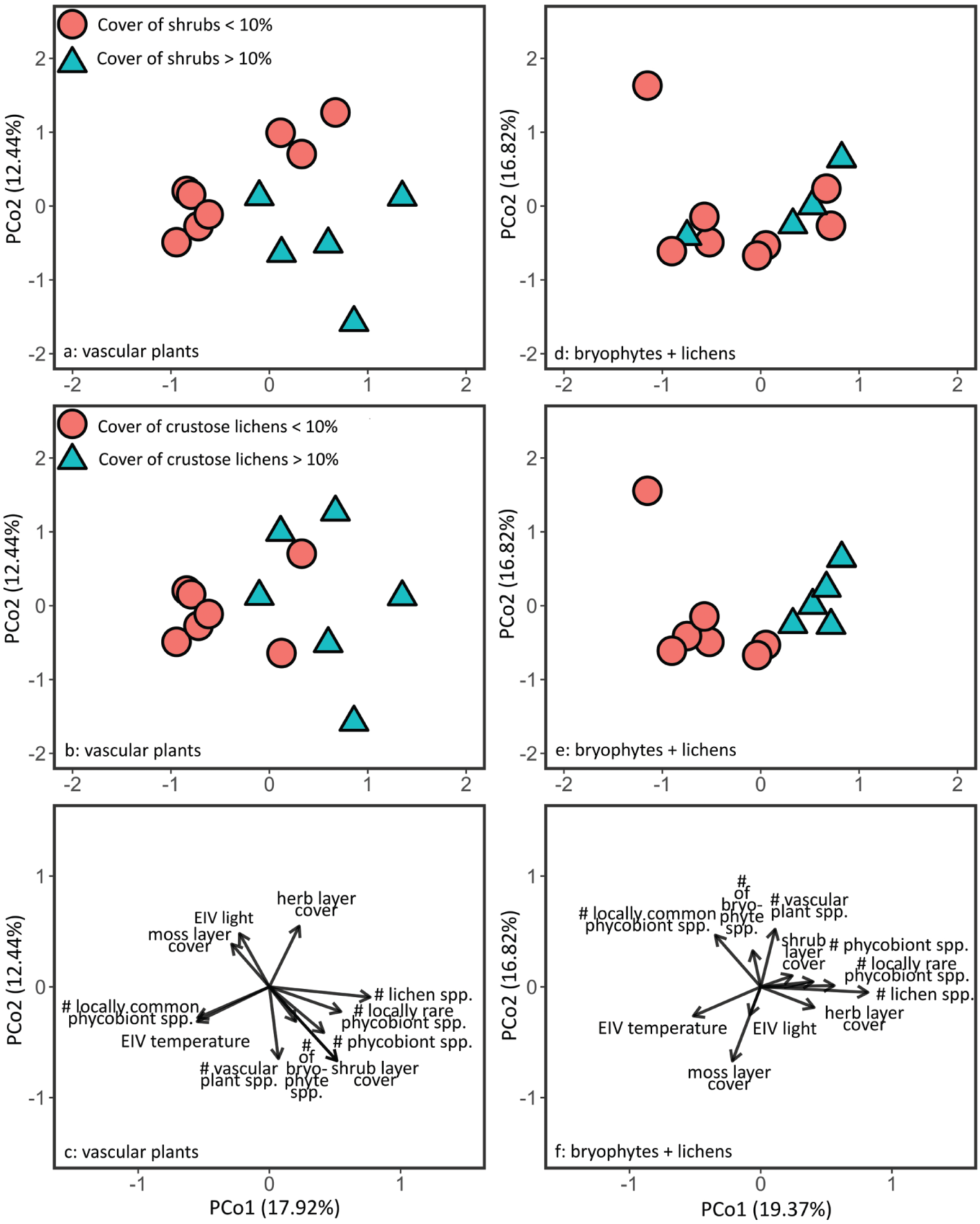
Ordination diagrams of principle coordinate analyses of community composition of vascular plants (a, b, c) and bryophytes + lichens (d, e, f). Colors and symbols indicate the proxy of successional development (in case of a and d it is the cover of shrubs being higher than 10%; in case of b and e it is the cover of crustose lichens being higher than 10%). Plots c and f show the passively superimposed environmental and biological variables. Ellenberg-type indicator values (EIVs) for light and temperature are calculated for vascular plants in case of plot c and for bryophytes in case of plot f. On axis labels the variation explained by particular ordination axis is given.

As indicated by the PCoA based on procrustes distances of taxa groups (Fig. 7), the variation of species composition of photobiont communities responded to the environment similarly as that of bryophyte communities, while clearly differing from vascular plants and lichens (Online Resource 1/Table S2). Note that the gradient of species diversity of taxa groups did not correspond to that of their community composition.

**Fig 7.**
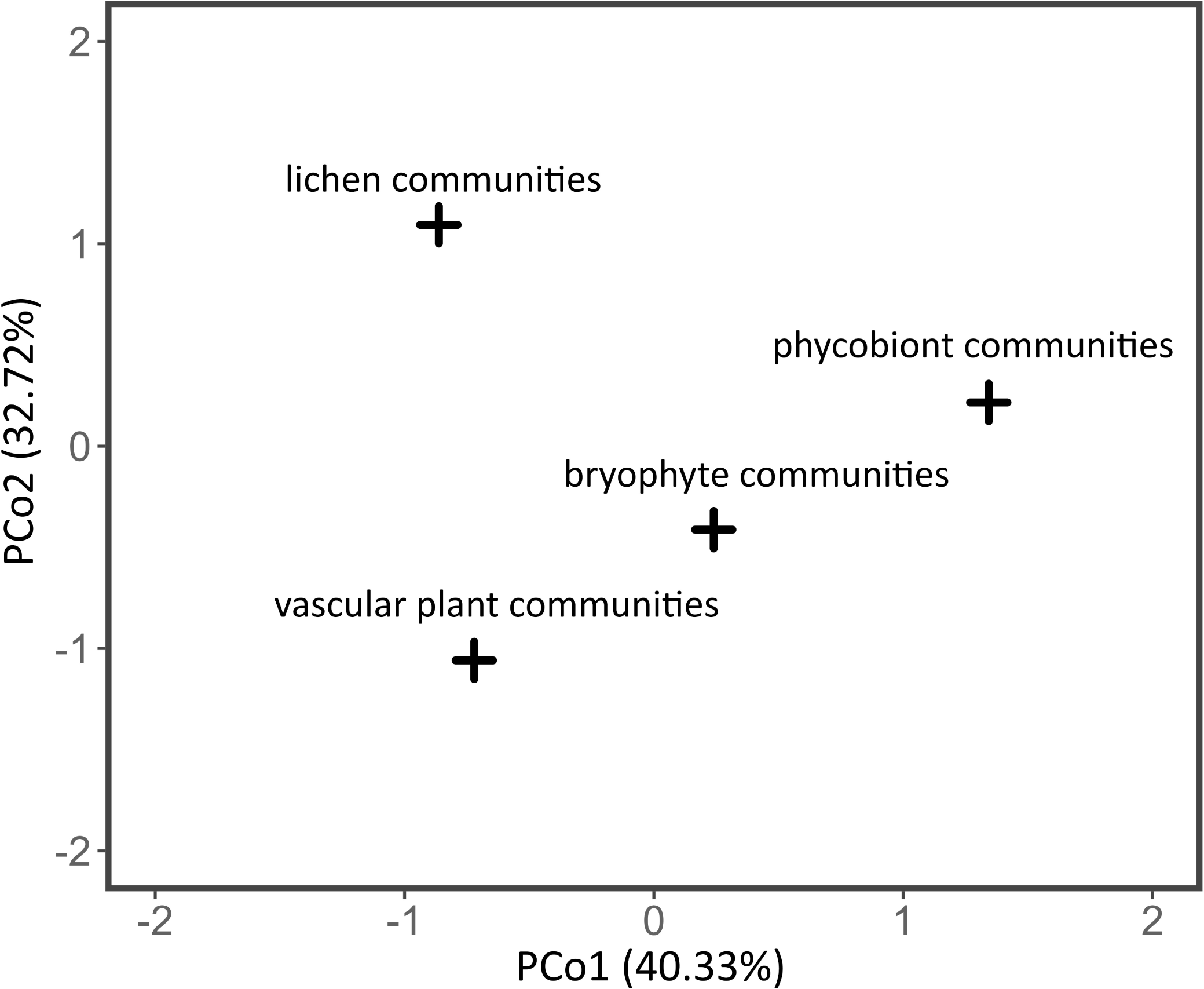
Ordination diagram (principle coordinate analysis) of procrustes distances between individual taxa groups. The diagram shows the difference in response of communities of individual taxa groups to ecological gradients covered by the sampled dataset. On axis labels the variation explained by particular ordination axis is given.

### Microalgal metabarcoding

Phycobiont diversity within particular lichen thalli (n=8) was inspected using Illumina metabarcoding. A total of 1,945,186 raw reads were generated, of which 1,240,063 passed the demultiplexing step and quality filter. This represented a mean of 155,007 (median 179,415) algal reads per sample, with a minimum of 38,329 and a maximum of 246,121 reads per sample. Filtered metabarcoding dataset consisted of 116 hits (4 to 44 per sample).

Abundances of recovered algal clades by sample are shown in Fig. 8. The predominant phycobiont covered from 52.4 to 98.8% reads. Thalli A523M, A554M, A563M, and A570M (assigned to the mycobiont OTU35) contained various *Asterochloris* species as the predominant phycobiont: lineage StA9 (sample A523M), *A. irregularis* (A563M) and *A. phycobiontica*/StA4/StA5 (A554M and A570M). The species-level lineages StA4, StA5 and *A. phycobiontica* are undistinguishable using ITS2 rDNA marker. Relative frequency of ASVs linked to *Asterochloris* spp. by sample is shown in Online Resource 2/Fig. S4. Thalli A597M, A598M, A633M, and A634M (assigned to mycobiont OTU2) contained predominant phycobiont from the trebouxiophycean lineage URa28.

**Fig. 8.**
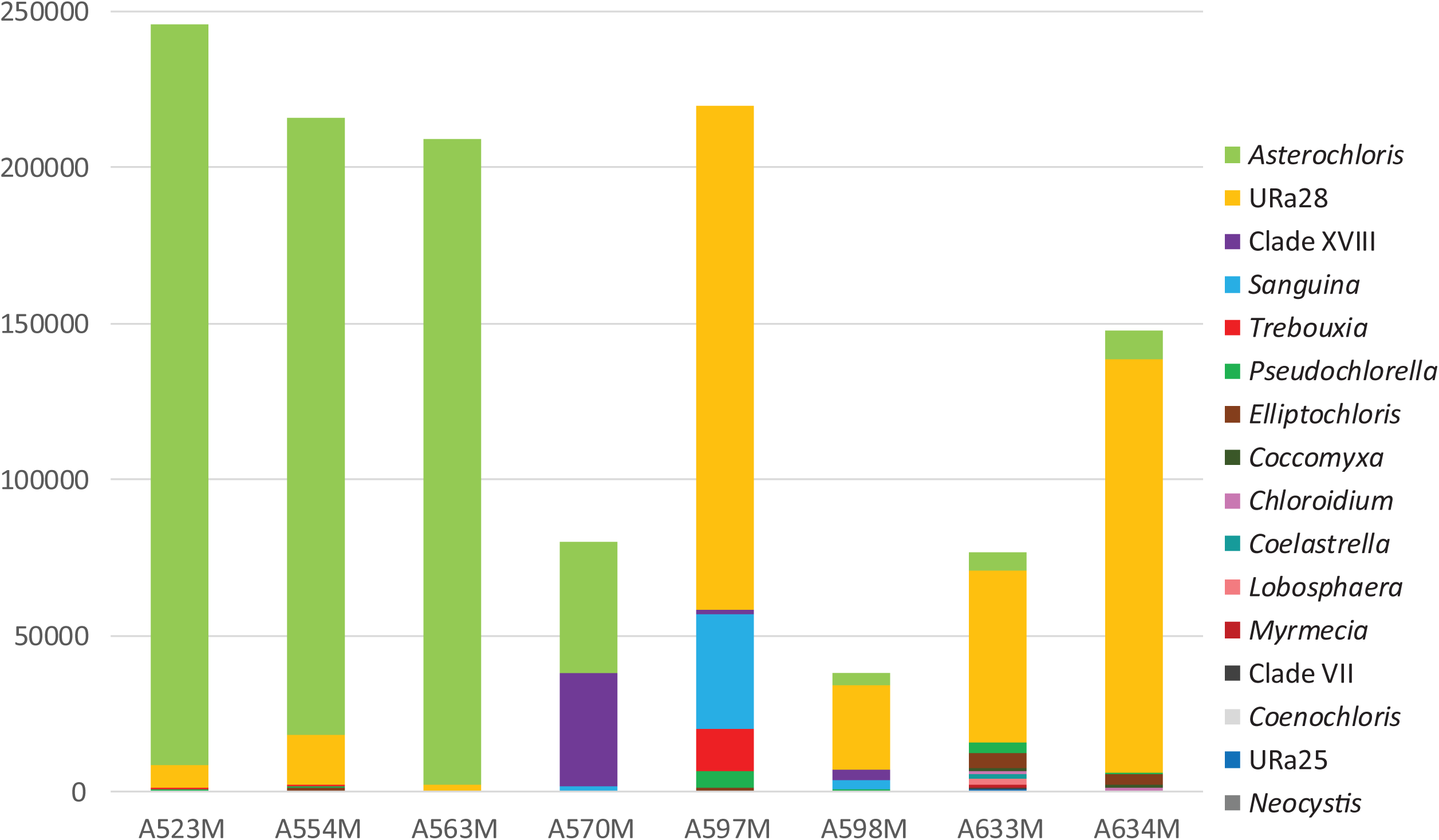
Eight *Stereocaulon* thalli selected for Illumina metabarcoding and abundances of recovered algal clades. A523M, A554M, A563M, A570M belong to mycobiont OTU35. A597M, A598M, A633M, A634M belong to mycobiont OTU2. Clade affiliations: URa25, URa28 *sensu* Ruprecht et al. (2014). Clades VII and XVIII were identified as new in present study. Amplicon sequence variants (ASVs) were sorted into these clades based on phylogenetic hypothesis presented in Online Resource 2/Fig. S3. Online Resource 1/Table S7 describes relative abundance of each ASV in each sample

### Diversity of soil algae

Phycobiont diversity in twelve soil samples was analyzed using Illumina metabarcoding. A total of 1,524,198 raw reads were generated, 876,596 of which passed the demultiplexing step and quality filter. This represented a mean of 73,049 (median 63,827) algal reads per sample, with a minimum of 18,255 and a maximum of 164,643 reads per sample. Filtered metabarcoding dataset consisted of 427 hits (4 to 89 per sample). The phylogenetic hypothesis resulting from Bayesian analysis of the ITS2 rDNA sequences obtained by Illumina metabarcoding of soil samples, selected lichen samples and reference sequences from GenBank is shown in Online Resource 2/Fig. S3. We recovered sequences in 44 well-supported clades, 27 of which contain exclusively soil algae, two contain exclusively phycobionts and 15 are shared by these two groups. Occurrence of particular clades of soil algae on each study plot is depicted in Online Resource 2/Fig. S5.

We also analyzed the occurrence of particular algal ITS2 haplotypes, including whole dataset obtained by Sanger sequencing (probably predominant phycobionts; Fig. 9a). Nine haplotypes were obtained by Sanger sequencing, in common with Illumina sequencing of soil and lichens. Vast majority of haplotypes was unique for soil (n=256) or Illumina lichen (n=79) datasets. On the contrary, 27 haplotypes were shared by soil and lichens but were not detected by Sanger sequencing of *Stereocaulon* in the study area. Same analysis restricted to haplotypes with frequency ≥1000 (in order to eliminate possible bias produced by errors from PCR and sequencing; Huse et al. 2010) showed a similar pattern (Fig. 9b).

**Fig. 9.**
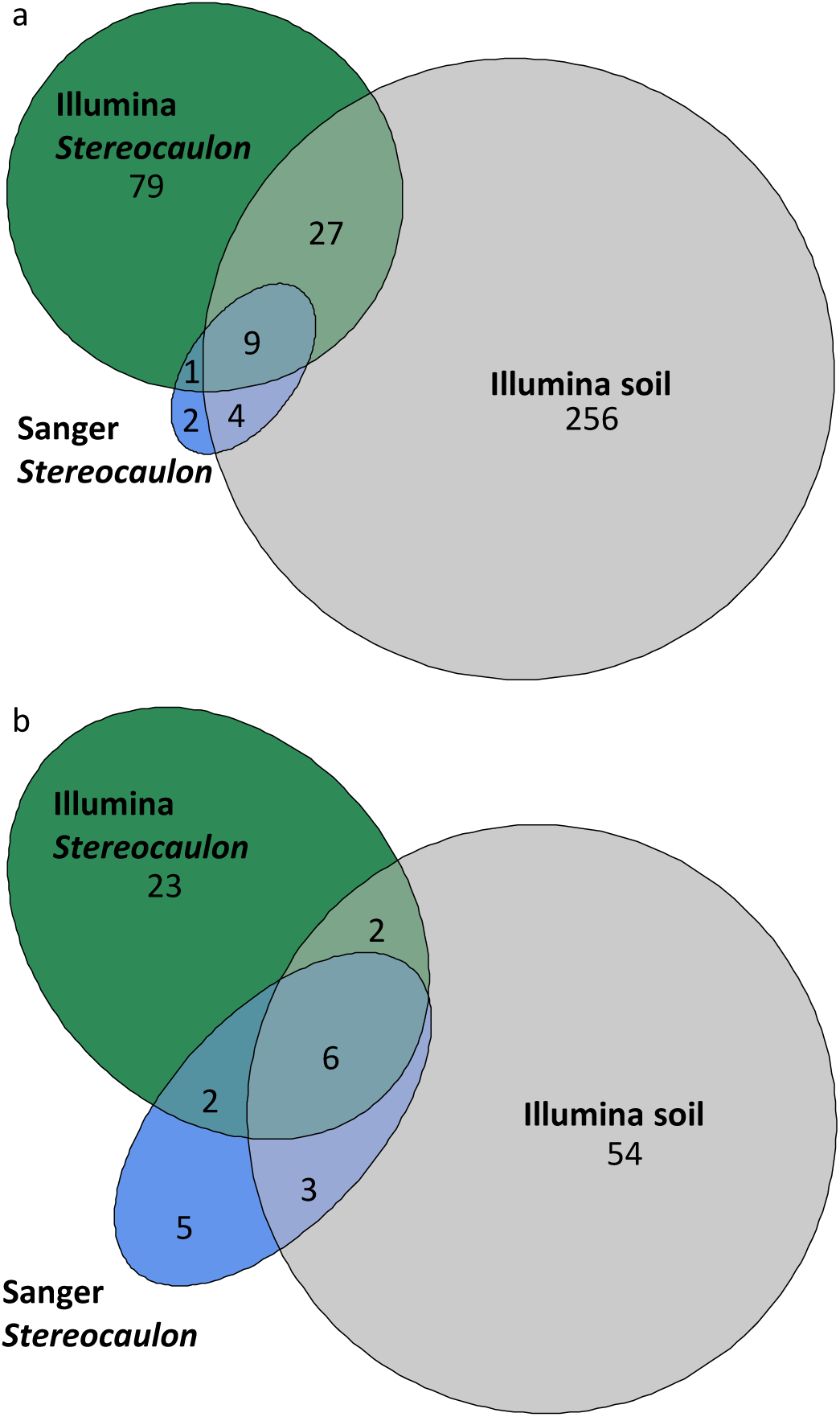
Euler diagrams depicting sets of algal ITS2 rDNA haplotypes recovered from selected *Stereocaulon* thalli (n=8) using Illumina metabarcoding, from all *Stereocaulon* samples (n=147) using Sanger sequencing and from soil samples (n=12) using Illumina metabarcoding. In case of Illumina metabarcoding sets, only haplotypes with frequency **a** ≥100 and **b** ≥1000 were included

## Discussion

### Change of community structure along a succession gradient

The succession on river gravel bars is an important driver of both species’ composition and diversity. It is well documented in case of vascular plants (e. g., Gilvear et al. 2008; Prach et al. 2014), but also applies to the microorganisms, such as soil bacteria or mycorrhizal fungi (Li et al. 2014; Sheng et al. 2017). However, different taxa groups response differently to this gradient. For example, vascular plants are known to follow a nested-community pattern when the highest species diversity is connected to the early-to mid-successional stages and the community is being impoverished with the ongoing succession (e. g., Walker and del Moral 2003; Corenblit et al. 2009; Chytrý et al. 2015). This corresponds to the pattern of vascular-plant-species-richness observed within this study.

On the other hand, the pattern observed for the phycobiont communities of *Stereocaluon* lichens differed. The phycobiont communities of early-vegetation stages were composed of relatively few species lineages such as *Asterochloris phycobiontica* or *Asterochloris* StA5 (Fig. 3), which are alpine and psychrophilic (Peksa and Škaloud 2011; Vančurová et al. 2018). As locally adapted, they are probably frequent also in source populations in the surroundings of the study plots. Therefore, newly emerged river gravel bars are easily colonized by *A. phycobiontica* and/or *Asterochloris* StA5 lineage. In subsequent stages, the observed species richness of phycobiont algae was mostly higher and these two species-level lineages were gradually substituted by others (Fig. 5). We hypothesize that some of them could be specialized to slightly different microhabitat conditions within particular plots. For example, clades 8, 12 and StA3 tolerate higher pH (Piercey-Normore and DePriest 2001; Bačkor et al. 2010; Vančurová et al. 2018; Steinová et al. 2019). On the river gravel bars of glacial floodplains, organisms with various substrate optima could coexist together because of heterogeneity of substrate transported by a river or glacier from more distant localities and various substrate layers. In the study area, acidophilic vascular plants and bryophytes dominated. However, occurrence of basophilic species such as *Didymodon fallax, Lophozia excisa, Syntrichia ruralis*, and *Veronica fruticans* is considered as indication of basic fractions in substrate.

The species richness of terricolous lichens on glacier forelands in Alps were positively correlated with time since deglaciation (Nascimbene et al. 2017) analogically to lichen species richness on deglaciated plots in maritime Antarctica (Favero-Longo et al. 2012). In both cases, most of species, once established, persisted to the oldest successional stages. On the river gravel bars the species richness of phycobionts was positively correlated with that of lichens (Fig. 4) but less with that of vascular plants and bryophytes (Fig. 6). This correlation probably indicates colonization of the localities by phycobionts and mycobionts using the similar dispersal strategies. On the other hand, we observed that the community composition variation (summarized by the procrustes tests; Fig. 7) of lichens and phycobiont communities of *Stereocaulon* lichens differed.

We hypothesized ongoing colonization of river gravel bars by the phycobionts, or alternatively, the acquisition of the phycobionts from soil *in situ* (Dal Grande et al. 2012; Fontaine and Beck 2012; Ohmura et al. 2019). We found several algal clades both in soil and lichens (Online Resource 2/Fig. S3), but the phycobiont pool seems to be independent of soil algae (Fig. 10). For example, the most frequent ITS2 rDNA haplotype among Sanger sequences (*A. phycobiontica*/StA4/StA5 recovered in 69% of all samples) was present only in two soil samples in a very low amount (212 and 135 reads, which is 2.4% and 3.1% of algal reads, respectively). This result supports our hypothesis that phycobionts originating from the surroundings colonize the recently emerged plots without substantial contribution of “soil seed bank”. Still, these results should be perceived as the basis for future research. The number of soil samples is rather limited, and some taxa could be undetected (Rippin et al. 2018). However, the taxonomic composition of algae occurring on river gravel bars is comparable to the pool of soil algae detected in the forefield of Damma glacier in the Swiss Alps (Frey et al. 2013). Notably, a significantly different algal community was found in an early-successional stage, which is in their case represented by a bare soil near to a receding glacier. The lichen phycobionts, including *Asterochloris*, were reported from soil in transitional and developed stage. Besides others, they found the lineage StA9, firstly reported as lichen phycobiont in the present study.

**Fig. 10.**
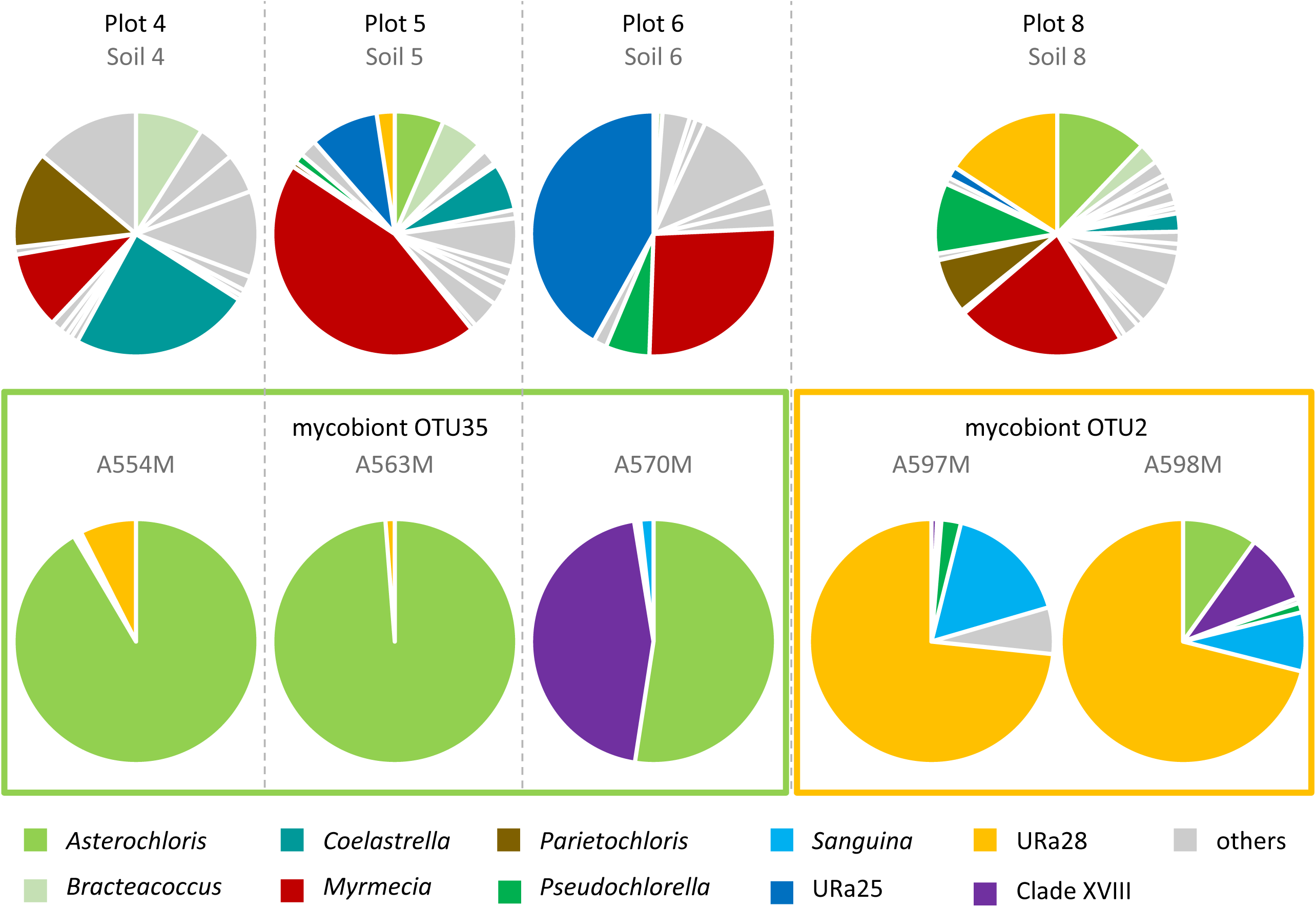
Relative abundances of algal clades within soil (first line) and *Stereocaulon* (second line) samples. Solely plots 4, 5, 6, and 8 with soil samples generating >5000 algal reads and with *Stereocaulon* samples analyzed using Illumina metabarcoding were displayed. Clade affiliations: URa25, URa28 *sensu* Ruprecht et al. (2014). Clade XVIII was identified as new in present study. Amplicon sequence variants (ASVs) were sorted into these clades based on phylogenetic hypothesis presented in Online Resource 2/Fig. S3

Beck et al. (2019) found two haplotypes *Stichococcus antarcticus*, phycobiont of *Placopsis* lichen in maritime Antarctica, exclusively in areas deglaciated for a long time with more developed soil and lichen community, unlike to other haplotypes occurring in whole area. The succession of vegetation causes numerous physical and chemical changes in soil. Therefore, the correlation between species richness of various organisms and successional stage (or age of the sample site) is frequently connected with changing soil characteristics, such as nutrient concentrations (Li et al. 2014; Sheng et al. 2017). However, according the Ellenberg indication values (Online Resource 1/Table S1), we did not detect such gradients along successional gradient in the study area. Contrarily, the gradients of distance from a river and the elevation above the water surface, are detectable in our data. Since the succession is blocked by river on gravel bars; succession gradient and distance from the river are naturally correlated. Nevertheless, no algae known from aquatic or humid environment (e.g. *Chloroidium*; Vančurová et al. 2018) were found as predominant phycobiont in samples growing in proximity to a river. Probably, the river disturbs the community occasionally but does not favor individuals tolerating submersion.

### Low specificity as an adaptive strategy

The specificity (i. e., the taxonomic range of acceptable partners; Rambold et al. 1998; Yahr et al. 2004, 2006) of both mycobiont and phycobiont has been considered as crucial characteristic of lichen interactions. The lower specificity towards symbiotic partners has been frequently reported as advantageous strategy in harsh environments (Romeike et al. 2002; Engelen et al. 2010), because it enables association with phycobionts adapted to a variety of conditions.

Two species-level lineages of *Stereocaulon* were recorded in the study area (Fig. 2). Both lineages are morphologically identical, also indistinguishable in the field. Overwhelming majority of samples belonged to OTU35, which has been reported to be low-specific towards its phycobionts (Vančurová et al. 2018). Frequently, the mycobionts belonging to this lineage adopted algae known generally as phycobionts of *Lepraria* lichens (*A. phycobiontica, A. echinata, A. leprarii*; Škaloud et al. 2015), in the study area. We hypothesize that such a low specificity (OTU35 associated with 13 species-level lineages of *Asterochloris* and in two cases also with representatives of other Trebouxiophacean phycobionts in study area) could facilitate colonizing heterogeneous and harsh habitat of river gravel bars of glacial floodplains.

On the other hand, none of the samples of *S. alpinum* OTU2 associated with the same phycobionts as the mycobiont OTU35, despite growing in the same environment. The OTU2 mycobiont mostly associated with trebouxiophycean alga URa28 as a phycobiont. This alga was previously detected in a soil sample from Canada either as a soil alga or possibly as a phycobiont of *Stereocaulon* sp. which was recorded in the same place (Hartmann et al. 2009).

### Algal plurality

The occurrence of more phycobiont species in a single lichen thallus (i.e., algal plurality) is a common but overlooked phenomenon (e. g. Bačkor et al. 2010; Moya et al. 2017; Onuț-Brännström et al. 2018) known also in *Stereocaulon* (Vančurová et al. 2018). However, various lichen species differ in the prevalence of algal plurality. Dal Grande et al. (2017) proved an occurrence of more phycobionts in 49.2% of *Lasallia hispanica* thalli but only in 1.7% of *L. pustulata* thalli. The algal plurality is mostly undetectable by the Sanger sequencing. Using the Illumina metabarcoding, we found more than one phycobiont in all selected samples from both mycobiont species-level lineages (Fig. 8). However, these samples were selected on the ground of difficulties with Sanger sequencing which could indicate the algal plurality (Paul et al. 2018). Illumina metabarcoding as well as the Sanger sequencing uncovered URa28 as a predominant phycobiont of *Stereocaulon alpinum* OTU2. In most cases, the phycobiont determined by the Sanger sequencing corresponds with a predominant phycobiont according to the high-throughput sequencing (Molins et al. 2018; Paul et al. 2018).

Even though the two mycobiont species-level lineages OTU2 and OTU35 differ in predominant phycobiont pool, they share the pool of other intrathalline algae, unlike two *Circinaria* spp. collected at the same location shared the predominant phycobiont, but showed a completely different pool of other intrathalline algae (Molins et al. 2018). The *Stereocaulon* OTUs differ significantly in the frequency of intrathalline algae (Fig. 8). Above all, most of the OTU35 (with *Asterochloris* as the predominant phycobiont) thalli contain also a small amount of URa28 algae and *vice versa*. Several algal clades interacted exclusively with one mycobiont species-level lineage, but their frequency was generally low. A comparable phycobiont pair *Trebouxia jamesii*/*Trebouxia* sp. TR9 found in *Ramalina farinacea* lichen is assumed to bring physiological benefits to that symbiotic system (Casano et al. 2011; Centeno et al. 2016). Alternatively, the minority phycobionts could occur in thalli without direct effects to the lichen. They might be used as a source of algal symbionts for other lichens in the locality.

## Conclusions

The diversity of phycobionts shifts along the succession gradient on the river gravel bars. We revealed positive correlation between the species richness of phycobionts and the species richness of the accompanying lichens in the locality indicating ongoing colonization of this habitat by both groups. The phycobiont communities of early-successional stages comprised relatively few species lineages. In subsequent stages, the observed species richness of phycobionts was mostly higher and the species-level lineages typical for early-successional stages were gradually substituted by others probably adapted to heterogenous microhabitat conditions of the river gravel bars.

A substantial phycobiont diversity (including 14 *Asterochloris* species-level lineages and three additional trebouxiophycean algae) recovered on the river gravel bars suggested low specificity of *Stereocaulon* mycobionts. This range of phycobionts could help to cope with heterogenous and dynamic conditions on river gravel bars of glacial floodplains.

We found more than one phycobiont in all selected samples belonging to both mycobiont OTUs (OTU2 and OTU35). *Asterochloris* phycobionts were recovered as the predominant phycobionts of OTU35 and trebouxiophycean lineage URa28 was the predominant phycobiont of OTU2. Although, broad community of other intrathalline algae was shared by both mycobionts.

This study provides novel insights into community structure of symbiotic microorganisms under harsh and dynamic conditions of river gravel bars. Additionally, it poses several challenging questions concerning cryptic lichen species and specificity towards the photobionts, dispersal of microscopic symbionts, an ecological function of additional intrathalline algae and an observed discrepancy between communities of soil and lichen algae.

## Supporting information

Online Resource 1

Online Resource 2

## Acknowledgements

We thank Helmut Mayrhofer and Jan Vondrák for their help with identification of a few critical lichen samples, Svatava Kubešová for help with bryophyte identification and Vít Grulich and Jiří Danihelka for identification of some specimens of vascular plants. This work was supported by the Charles University Science Foundation project GAUK 946417, the Primus Research Programme of Charles University no. SCI/13 and by the long-term research development project RVO 67985939.

## Supplementary material

**Table S1** Location, environmental and vegetation characteristics of study plots

**Table S2** Species composition of gravel bar vegetation plots. Species are ranked by decreasing frequency of occurrence. The abundance of species is given on the nine-degree Braun-Blanquet scale. All relevés are stored in Gravel bar vegetation database (Kalníková and Kudrnovsky 2017)

**Table S3** Primers used in this study

**Table S4** GenBank accession numbers, phycobiont species-level lineage, mycobiont OTU, substrate and affiliation to study plot of *Stereocaulon* samples

**Table S5** Substitution models selected for each partition of *Asterochloris* and *Stereocaulon* (mycobiont) datasets and for algal ITS2 rDNA dataset using the Bayesian information criterion (BIC) as implemented in JModelTest2 (Guindon & Gascuel 2003, Darriba et al. 2012)

**Table S6** Accession numbers of *Asterochloris* reference sequences retrieved from GenBank

**Table S7** Relative abundance of each ASV in each *Stereocaulon* sample

**Fig. S1** Examples of study plots belonging to different succession stages: **a, b** succession stage 1 (plots 5 and 9), **c** succession stage 2 (plot 2), **d** succession stage 3 (plot 3)

**Fig. S2** Rarefaction curves for 13 study plots. Vertical lines are drawn at sample size of n=5 and sample size of n=10

**Fig. S3** Phylogenetic hypothesis resulting from Bayesian analysis of the algal ITS2 rDNA sequences obtained by Illumina metabarcoding of soil samples, selected lichen samples and reference sequences from GenBank. Clade affiliations: URa25, URa28 *sensu* Ruprecht et al. (2014). Clades I– XVIII were identified as new in present study. Values at the nodes indicate the statistical supports of Bayesian posterior probability

**Fig. S4** Relative frequency of ASVs linked to various *Asterochloris* species. Solely samples with *Asterochloris* as a predominant phycobiont (>50% algal reads belonged to *Asterochloris*) were displayed

**Fig. S5** Frequency of ASVs linked to algal clades recovered from soil samples. Clade affiliations: URa25, URa28 *sensu* Ruprecht et al. (2014). Clades I–XVIII were identified as new in present study. Amplicon sequence variants (ASVs) were sorted into these clades based on phylogenetic hypothesis presented in Online Resource 2/Fig. S3

