## Supplementary material for "Symbiosis between river and dry lands: phycobiont dynamics on river gravel bars": Online Resource 1

**List of contents:**

Table S1 Page 2

Table S2 Page 4

Table S3 Page 7

Table S4 Page 8

Table S5 Page 13

Table S6 Page 14

Table S7 Page 15

**Table S1** Location, environmental and vegetation characteristics of study plots

| Plot number | Succession stage | Locality | Altitude (m) | Cover total (%) | Cover tree layer (%) | Cover shrub layer (%) | Cover herb layer (%) | Cover moss layer (%) | Cover lichen layer (%) | Cover crustose lichens (%) | Cover Stereocaulon (%) | River distance (m) | Height above river (m) | No of. vascular plants |
|---|---|---|---|---|---|---|---|---|---|---|---|---|---|---|
| 1 | 1 | Morteratsch | 2070 | 70 | 0 | 0 | 25 | 55 | 15 | 0.0 | 8 | 10.0 | 0.8 | 13 |
| 2 | 2 | Morteratsch | 2018 | 80 | 0 | 35 | 15 | 25 | 5 | 3.0 | 3 | 35.0 | 2.5 | 23 |
| 3 | 3 | Morteratsch | 2026 | 85 | 10 | 30 | 10 | 20 | 15 | 16.6 | 4 | 240.0 | 13.0 | 27 |
| 4 | 1 | RosegI | 2034 | 10 | 0 | 0 | 8 | 2 | 1 | 1.0 | 2 | 15.0 | 0.5 | 25 |
| 5 | 1 | RosegI | 2031 | 50 | 0 | 0 | 15 | 30 | 10 | 17.5 | 8 | 2.0 | 0.7 | 13 |
| 6 | 2 | RosegI | 2040 | 50 | 0 | 5 | 15 | 33 | 3 | 7.8 | 2 | 75.0 | 1.0 | 22 |
| 7 | 3 | RosegI | 2050 | 70 | 0 | 15 | 20 | 35 | 5 | 10.5 | 3 | 20.0 | 2.5 | 22 |
| 8 | 2 | RosegII | 2012 | 90 | 0 | 5 | 10 | 80 | 5 | 4.0 | 3 | 10.0 | 1.5 | 18 |
| 9 | 1 | RosegII | 1997 | 70 | 0 | 0 | 5 | 70 | 2 | 7.7 | 2 | 2.5 | 0.4 | 10 |
| 10 | 1 | Lonza | 1995 | 70 | 0 | 0 | 30 | 45 | 7 | 16.6 | 3 | 0.3 | 1.0 | 17 |
| 11 | 1 | Lonza | 2027 | 65 | 0 | 0 | 15 | 55 | 2 | 4.9 | 2 | 4.0 | 1.0 | 13 |
| 12 | 2 | Lonza | 2003 | 80 | 0 | 15 | 15 | 40 | 10 | 17.5 | 8 | 16.0 | 0.9 | 24 |
| 13 | 3 | Lonza | 2007 | 60 | 5 | 20 | 15 | 25 | 5 | 18.3 | 3 | 200.0 | 10.0 | 17 |

| | No. of bryophytes | No of. lichens | EIV light vascular plants | EIV temperature vascular plants | EIV continentality vascular plants | EIV humidity vascular plants | EIV reaction vascular plants | EIV nutrients vascular plants | EIV light bryophytes | EIV temperature bryophytes | EIV continentality bryophytes |
|---|---|---|---|---|---|---|---|---|---|---|---|
| plot1 | 4 | 6 | 8.00 | 2.89 | 3.73 | 4.55 | 5.78 | 2.70 | 8.67 | 2.50 | 5.50 |
| plot2 | 7 | 11 | 7.47 | 3.33 | 4.11 | 5.00 | 5.75 | 3.00 | 7.83 | 2.25 | 5.50 |
| plot3 | 7 | 21 | 7.50 | 3.13 | 4.20 | 4.82 | 5.28 | 2.91 | 7.75 | 2.60 | 5.75 |
| plot4 | 5 | 3 | 7.75 | 3.21 | 4.35 | 4.70 | 6.38 | 3.32 | 8.00 | 2.50 | 5.67 |
| plot5 | 2 | 20 | 7.82 | 3.30 | 3.91 | 4.82 | 5.75 | 3.27 | 9.00 | 2.50 | 5.50 |
| plot6 | 2 | 9 | 7.71 | 3.50 | 4.06 | 5.00 | 6.14 | 3.25 | 9.00 | 2.50 | 5.50 |
| plot7 | 4 | 14 | 7.80 | 3.00 | 4.00 | 4.85 | 5.93 | 2.74 | 7.50 | 2.50 | 5.50 |
| plot8 | 3 | 10 | 7.81 | 3.55 | 4.12 | 4.88 | 6.13 | 3.00 | 8.33 | 2.50 | 6.00 |
| plot9 | 2 | 11 | 7.86 | 4.00 | 4.13 | 4.13 | 5.83 | 3.00 | 9.00 | 2.50 | 5.50 |
| plot10 | 5 | 16 | 7.67 | 2.75 | 3.86 | 4.67 | 4.91 | 2.80 | 7.80 | 2.50 | 5.33 |
| plot11 | 2 | 8 | 7.91 | 2.78 | 4.50 | 5.33 | 6.00 | 3.55 | 9.00 | 2.50 | 5.50 |
| plot12 | 3 | 18 | 7.89 | 3.29 | 4.06 | 4.50 | 5.92 | 2.83 | 7.67 | 2.50 | 5.67 |
| plot13 | 5 | 15 | 7.86 | 2.69 | 4.15 | 4.46 | 4.44 | 2.62 | 8.50 | 2.67 | 5.50 |

**Table S1** cont.

| | EIV humidity bryophytes | EIV reaction bryophytes | Phycobiont species (sample size 5) | Phycobiont species (sample size 10) | GPS coordinates |
|---|---|---|---|---|---|
| plot1 | 1.67 | 4.00 | 2.28 | 3.00 | 46.4308528, 9.9357028 |
| plot2 | 4.00 | 4.57 | 3.09 | 4.73 | 46.4305556; 9.9350000 |
| plot3 | 5.20 | 3.80 | 3.50 | 4.97 | 46.4332500, 9.9332500 |
| plot4 | 2.25 | 5.75 | 2.36 | 2.97 | 46.4253611, 9.8604444 |
| plot5 | 1.50 | 4.00 | 3.66 | 5.30 | 46.4250278, 9.8602500 |
| plot6 | 1.50 | 4.00 | 2.68 | 4.08 | 46.4238333, 9.8590278 |
| plot7 | 2.75 | 5.00 | 2.78 | 4.00 | 46.4215556, 9.8586111 |
| plot8 | 2.33 | 4.50 | 2.73 | 3.00 | 46.4341111, 9.8649722 |
| plot9 | 1.50 | 4.00 | 2.95 | 4.00 | 46.4376944, 9.8702778 |
| plot10 | 2.80 | 5.25 | 2.41 | 2.91 | 46.4459722, 7.8999167 |
| plot11 | 1.50 | 4.00 | 2.00 | NA | 46.4473056, 7.9040556 |
| plot12 | 2.33 | 5.33 | 3.39 | 5.00 | 46.4463333, 7.9000833 |
| plot13 | 2.25 | 5.67 | 2.91 | 4.00 | 46.4465000, 7.8997778 |

**Table S2** Species composition of gravel bar vegetation plots. Species are ranked by decreasing frequency of occurrence. The abundance of species is given on the nine-degree Braun-Blanquet scale. All relevés are stored in Gravel bar vegetation database (Kalníková and Kudrnovsky 2017)

| **Succession stage** | **1** | **2** | **3** |
|---|---|---|---|
| **Plot number** | 11 | 1 | 1 |
| | 145901 | 2682 | 373 |
| **Tree layer** | | | |
| *Larix decidua* | ...... | .... | a.1 |
| **Shrub layer** | | | |
| *Salix helvetica* | ...... | ++.a | a.b |
| *Salix daphnoides* | ...... | b+.. | .+. |
| *Salix myrsinifolia* | ...... | 1... | 1.. |
| *Salix species* | ...... | +... | ..+ |
| *Juniperus communis* subp. *nana* | ...... | .... | .1+ |
| *Salix purpurea* | ...... | 1... | ... |
| *Alnus viridis* | ...... | .1.. | ... |
| *Myricaria germanica* | ...... | ..m. | ... |
| *Rhododendron ferrugineum* | ...... | .... | 1.. |
| *Salix appendiculata* | ...... | .... | +.. |
| *Pinus cembra* | ...... | .... | +.. |
| *Salix foetida* | ...... | .... | .1. |
| *Larix decidua* | ...... | .... | .+. |
| *Picea abies* | ...... | .... | ..+ |
| **Tree or shrub juveniles in the herb layer** | | | |
| *Populus tremula* | ...... | +... | ... |
| *Salix species* | .+.... | .... | ... |
| *Pinus cembra* | ...... | .... | r.. |
| *Larix decidua* | ...... | .... | +.+ |
| *Myricaria germanica* | .+.... | .... | ... |
| **Herb layer** | | | |
| *Epilobium fleischeri* | b+a1aa | aaa1 | +m+ |
| *Tolpis staticifolia* | 1.m+m1 | 1111 | +a+ |
| *Poa alpina* | ++++++ | +1++ | +.+ |
| *Poa nemoralis* | +11++. | +m1+ | .m. |
| *Trifolium pallescens* | .++.11 | 1++a | .+a |
| *Sempervivum arachnoideum* | ++.+.. | ++++ | 1+. |
| *Saxifraga paniculata* | ++..1. | +1++ | .+. |
| *Anthyllis vulneraria* s. lat. | .a1+.. | .1m1 | ++. |
| *Achillea erba-rotta* subsp. *moschata* | ..+.++ | ..++ | +1+ |
| *Rumex scutatus* | ++.+.. | .1+. | .+. |
| *Trifolium badium* | +r.... | a++. | .+. |
| *Festuca rubra* agg. | .+.+.+ | .1+. | 1.. |
| *Saxifraga bryoides* | ++...+ | ++.. | ... |
| *Erigeron acris* subsp. *angulosus* | r++... | .r+. | ... |
| *Lotus alpinus* | ....++ | +..+ | +.. |
| *Campanula cochleariifolia* | ..+..r | .+.+ | ... |
| *Agrostis schraderiana* | .+...+ | .... | .+. |
| *Scorzoneroides autumnalis* | .+.... | .++. | ... |
| *Minuartia verna* | .+.... | .+.+ | ... |
| *Cerastium arvense* subsp. *strictum* | .r.... | ...+ | +.. |
| *Sempervivum montanum* | ..+.1. | .... | ..1 |
| *Veronica fruticans* | ....+. | .... | .++ |
| *Euphrasia species* | ...... | +..+ | +.. |
| *Juncus trifidus* | ...... | +..+ | ..r |
| *Luzula spicata* | ...... | ...+ | +.r |
| *Saxifraga aizoides* | +..... | +... | ... |
| *Biscutella laevigata* | .+.... | ..+. | ... |
| *Trifolium pratense* | ..+... | .... | 1.. |
| *Festuca species* | ....+. | ...+ | ... |
| *Leucanthemopsis alpina* | ....+. | .... | .r. |
| *Euphrasia minima* | ....+. | .... | ..+ |
| *Agrostis stolonifera* agg. | .....+ | ...+ | ... |

**Table S2** cont.

<table>
<tr><td>Pyrola minor</td><td>......|+...|+..</td></tr>
<tr><td>Galium species</td><td>......|.+.+|...</td></tr>
<tr><td>Parnassia palustris</td><td>......|.+..|r..</td></tr>
<tr><td>Pilosella officinarum</td><td>......|....|++.</td></tr>
<tr><td>Silene rupestris</td><td>+.....|....|...</td></tr>
<tr><td>Taraxacum species</td><td>r.....|....|...</td></tr>
<tr><td>Poa compressa</td><td>.+....|....|...</td></tr>
<tr><td>Taraxacum Sec. Alpina</td><td>.+....|....|...</td></tr>
<tr><td>Festuca quadriflora</td><td>.+....|....|...</td></tr>
<tr><td>Deschampsia cespitosa</td><td>.+....|....|...</td></tr>
<tr><td>Campanula species</td><td>.r....|....|...</td></tr>
<tr><td>Arabis species</td><td>.r....|....|...</td></tr>
<tr><td>Saxifraga species</td><td>..+...|....|...</td></tr>
<tr><td>Saxifraga oppositifolia</td><td>..+...|....|...</td></tr>
<tr><td>Poa species</td><td>...r..|....|...</td></tr>
<tr><td>Calamagrostis species</td><td>...r..|....|...</td></tr>
<tr><td>Jacobaea incana</td><td>....+.|....|...</td></tr>
<tr><td>Alchemilla species</td><td>....+.|....|...</td></tr>
<tr><td>Botrychium lunaria</td><td>....+.|....|...</td></tr>
<tr><td>Myosotis alpestris</td><td>....r.|....|...</td></tr>
<tr><td>Festuca rupicaprina</td><td>.....+|....|...</td></tr>
<tr><td>Sempervivum species</td><td>.....r|....|...</td></tr>
<tr><td>Hypericum species</td><td>......|+...|...</td></tr>
<tr><td>Agrostis rupestris</td><td>......|+...|...</td></tr>
<tr><td>Saxifraga aspera</td><td>......|+...|...</td></tr>
<tr><td>Hieracium species</td><td>......|.+..|...</td></tr>
<tr><td>Scorzoneroides helvetica</td><td>......|..+.|...</td></tr>
<tr><td>Tussilago farfara</td><td>......|..r.|...</td></tr>
<tr><td>Veronica species</td><td>......|...+|...</td></tr>
<tr><td>Sedum species</td><td>......|...+|...</td></tr>
<tr><td>Arabis caerulea</td><td>......|...r|...</td></tr>
<tr><td>Carex sempervirens</td><td>......|...r|...</td></tr>
<tr><td>Anthoxanthum alpinum</td><td>......|....|1..</td></tr>
<tr><td>Polystichum lonchitis</td><td>......|....|+..</td></tr>
<tr><td>Avenella flexuosa</td><td>......|....|+..</td></tr>
<tr><td>Thesium alpinum</td><td>......|....|r..</td></tr>
<tr><td>Bellardiochloa variegata</td><td>......|....|.1.</td></tr>
<tr><td>Thymus pulegioides agg.</td><td>......|....|.+.</td></tr>
<tr><td>Trifolium species</td><td>......|....|.+.</td></tr>
<tr><td>Carex atrata subsp. aterrima</td><td>......|....|.+.</td></tr>
<tr><td>Cardamine resedifolia</td><td>......|....|..+</td></tr>
<tr><td>Moss layer</td><td></td></tr>
<tr><td>Bryophytes</td><td></td></tr>
<tr><td>Racomitrium canescens</td><td>3+3533|3353|b3b</td></tr>
<tr><td>Polytrichum piliferum</td><td>1.++.+|+1.+|...</td></tr>
<tr><td>Polytrichum juniperinum</td><td>.r..1.|..+.|+1+</td></tr>
<tr><td>Ceratodon purpureus</td><td>+...+.|..+.|.++</td></tr>
<tr><td>Cephaloziella species</td><td>1.....|....|+.+</td></tr>
<tr><td>Bryum species</td><td>.+....|....|+..</td></tr>
<tr><td>Barbula unguiculata</td><td>.+....|....|...</td></tr>
<tr><td>Didymodon fallax</td><td>.+....|....|...</td></tr>
<tr><td>Syntrichia ruralis</td><td>....+.|....|...</td></tr>
<tr><td>Bryum capillare s. lat.</td><td>....+.|....|...</td></tr>
<tr><td>Bryum argenteum</td><td>......|+...|...</td></tr>
<tr><td>Bryum caespiticium</td><td>......|+...|...</td></tr>
<tr><td>Sanionia uncinata</td><td>......|+...|...</td></tr>
<tr><td>Bryoerythrophyllum recurvirostrum</td><td>......|+...|...</td></tr>
<tr><td>Pohlia filum</td><td>......|+...|...</td></tr>
<tr><td>Tortella tortuosa</td><td>......|...+|...</td></tr>
<tr><td>Aulacomnium palustre</td><td>......|....|1..</td></tr>
<tr><td>Lophozia excisa</td><td>......|....|+..</td></tr>
<tr><td>Oncophorus virens</td><td>......|....|+..</td></tr>
<tr><td>Scapania undulata</td><td>......|....|+..</td></tr>
<tr><td>Amblystegium serpens</td><td>......|....|.+.</td></tr>
<tr><td>Tortella inclinata</td><td>......|....|..+</td></tr>
</table>

**Table S2** cont.

| | | | |
|---|---|---|---|
| **Lichens** | | | |
| *Stereocaulon alpinum* | a+a+1+ | 1+1a | m11 |
| *Peltigera rufescens* | 1+1++. | r11+ | r+1 |
| *Peltigera didactyla* agg. | +.+rr. | r+++ | +m. |
| *Cladonia fimbriata* | +..r+. | +.r1 | 11+ |
| *Lecanora polytropa* | ..+r+r | .+.+ | +++ |
| *Acarospora cf. veronensis* | ..+.+r | .r++ | +++ |
| *Cladonia pyxidata* | r.+.++ | +..+ | +r. |
| *Rhizocarpon geographicum* | ..+.+r | .+.+ | +++ |
| *Caloplaca subpallida* | ..+r+. | ..r+ | r.+ |
| *Bellemerea sanguinea* | ..+.+r | .+.. | +++ |
| *Rhizocarpon polycarpum* | ..+.+. | .r.+ | +++ |
| *Placynthiella icmalea* | .r..r. | r..+ | ..+ |
| *Cladonia chlorophaea* agg. | ..+.+. | r... | r+. |
| *Physcia dubia* | ..+..r | .... | r++ |
| *Lecidea cervinicola* | ..r.r. | ...r | +.+ |
| *Cladonia rei* | r..... | ..r. | rr. |
| *Candelariella vitellina* | ..+.+. | .... | +.+ |
| *Myriolecis dispersa* agg. | ..r... | ...r | ..r |
| *Umbilicaria cylindrica* | ..+... | .+.. | ... |
| *Peltigera lepidophora* | ..r... | ..+. | ... |
| *Caloplaca sinapisperma* | ...r.. | ..r. | ... |
| *Staurothele areolata* | .....r | ...+ | ... |
| *Cladonia cariosa* | ...... | r.r. | ... |
| *Cladonia macrophyllodes* | ...... | r..+ | ... |
| *Bellemerea cinereorufescens* | ...... | ...+ | ..r |
| *Candelariella species* | ..+... | .... | ... |
| *Cetraria ericetorum* | ..+... | .... | ... |
| *Placynthiella species* | ..r... | .... | ... |
| *Trapeliopsis flexuosa* | ...r.. | .... | ... |
| *Rinodina laxa* | ...r.. | .... | ... |
| *Amandinea punctata* | ...r.. | .... | ... |
| *Agonimia vouauxii* | ...r.. | .... | ... |
| *Placynthiella oligotropha* | ....+. | .... | ... |
| *Caloplaca stillicidiorum* | ...... | r... | ... |
| *Bacidia bagliettoana* | ...... | r... | ... |
| *Sarcogyne privigna* | ...... | ...r | ... |
| *Peltigera elisabethae* | ...... | ...r | ... |
| *Peltigera canina* | ...... | .... | 1.. |
| *Immersaria athroocarpa* | ...... | .... | r.. |
| *Porpidia macrocarpa* | ...... | .... | r.. |
| *Anaptychia bryorum* | ...... | .... | r.. |
| *Xanthoria elegans* | ...... | .... | .r. |

**Table S3** Primers used in this study

| Name | Sequence | | Reference |
|---|---|---|---|
| **nr-SSU-1780-5’** | 5’-CTG CGG AAG GAT CAT TGA TTC-3’ | algal ITS region, algal-specific | Piercey-Normore & DePriest 2001 |
| **ITS1-F-5’** | 5’- CTT GGT CAT TTA GAG GAA GTA A -3’ | fungal ITS region, fungal-specific | Gardes & Bruns 1993 |
| **ITS4-3’** | 5’-TCC TCC GCT TAT TGA TAT GC-3’ | algal and fungal ITS region, universal | White et al. 1990 |
| **ActinF2 Astero-5’** | 5’-AGC GCG GGT ACA GCT TCA C-3’ | actin type I locus, algal specific | Škaloud & Peksa, 2010 |
| **ActinR2 Astero-3’** | 5’-CAG CAC TTC AGG GCA GCG GAA-3’ | actin type I locus, algal specific | Škaloud & Peksa, 2010 |
| **1378-Chlorophyta** | 5’-TTG CCT TGT CAG GTT GAT TCC GG-3‘ | Illumina sequencing of ITS2 | this study |
| **5.8F-Chlorophyta** | 5’-GAA TTC CGT GAA CCA TCG AAT CTT T-3‘ | Illumina sequencing of ITS2 | this study |

**Table S4** GenBank accession numbers, phycobiont species-level lineage, mycobiont OTU, substrate and affiliation to study plot of *Stereocaulon* samples

| algal ITS accession | fungal ITS accession | actin accession | sample ID | phycobiont | mycobiont | | | substrate/surroundings | | | | |
|---|---|---|---|---|---|---|---|---|---|---|---|---|
| | | | | | | gravel | soil | moss | stone | needles | tree/shrub | plants |
| **study plot 1** | | | | | | | | | | | | |
| | | | VancurovaA509 | *Asterochloris StA5* | OTU35 | x | | | | | | |
| | | | VancurovaA510 | *Asterochloris StA5* | OTU35 | x | | | | | | |
| | | | VancurovaA511 | *Asterochloris StA5* | OTU35 | x | | | | | | |
| | | | VancurovaA512 | *Asterochloris StA5* | OTU35 | | x | x | | | | |
| | | | VancurovaA513 | *Asterochloris StA5* | OTU35 | | x | x | | | | |
| | | | VancurovaA514 | *Asterochloris phycobiontica* | OTU35 | | x | x | | | | |
| | | | VancurovaA515 | *Asterochloris phycobiontica* | OTU35 | | x | x | | | | |
| | | | VancurovaA516 | *Asterochloris clade12* | OTU35 | | x | x | | | | |
| | | | VancurovaA528 | *Asterochloris StA5* | OTU35 | x | | | | | | |
| | | | VancurovaA529 | *Asterochloris StA5* | OTU35 | x | | | | | | |
| **study plot 2** | | | | | | | | | | | | |
| | | | VancurovaA517 | *Asterochloris StA5* | OTU35 | x | | | | | | |
| | | | VancurovaA518 | *Asterochloris clade12* | OTU35 | x | | | | | | |
| | | | VancurovaA519 | *Asterochloris glomerata* | OTU35 | | | x | | | | |
| | | | VancurovaA520 | *Asterochloris StA5* | OTU35 | | | x | | | | |
| | | | VancurovaA521 | *Asterochloris clade8* | OTU35 | | | | x | | | |
| | | | VancurovaA522 | *Asterochloris StA5* | OTU35 | | x | | | | x | |
| | | | VancurovaA523V | *Asterochloris StA9* | OTU35 | | | | | | | |
| | | | VancurovaA524 | *Asterochloris StA5* | OTU35 | | | | x | | | |
| | | | VancurovaA525 | *Asterochloris StA5* | OTU35 | | | | x | | | |
| | | | VancurovaA526 | *Asterochloris StA9* | OTU35 | | x | | | | | |
| | | | VancurovaA527 | *Asterochloris StA5* | OTU35 | x | x | | | | | |
| **study plot 3** | | | | | | | | | | | | |
| | | | VancurovaA530 | *Asterochloris aff. italiana* | OTU35 | | | | x | | | |
| | | | VancurovaA531 | *Asterochloris leprarii* | OTU35 | | | | | x | x | |
| | | | VancurovaA532 | *Asterochloris phycobiontica* | OTU35 | | | x | | | | x |
| | | | VancurovaA533 | *Asterochloris StA5* | OTU35 | x | | x | | | | |
| | | | VancurovaA534 | *Asterochloris StA5* | OTU35 | | | x | | x | | |
| | | | VancurovaA535 | *Asterochloris phycobiontica* | OTU35 | | | x | | | x | |
| | | | VancurovaA536 | *Asterochloris phycobiontica* | OTU35 | | | x | | | | |
| | | | VancurovaA537 | *Asterochloris aff. italiana* | OTU35 | x | | x | | | | |
| | | | VancurovaA538 | *Asterochloris StA5* | OTU35 | | | x | | x | x | |
| | | | VancurovaA539 | *Asterochloris phycobiontica* | OTU35 | x | | x | | | | |
| | | | VancurovaA540 | *Asterochloris phycobiontica* | OTU35 | | | | | x | x | |
| | | | VancurovaA541 | *Asterochloris lobophora* | OTU35 | | x | | | | x | |
| | | | VancurovaA542 | *Asterochloris StA5* | OTU35 | x | | x | | | | |

**Table S4** cont.

| | | | | | | | | substrate/surroundings | | | |
|---|---|---|---|---|---|---|---|---|---|---|---|
| **algal ITS accession** | **fungal ITS accession** | **actin accession** | **sample ID** | **phycobiont** | **mycobiont** | **gravel** | **soil** | **moss** | **stone** | **needles** | **tree/shrub plants** |
| | | | VancurovaA543 | *Asterochloris StA4* | OTU35 | | | | x | | |
| | | | VancurovaA544 | *Asterochloris aff. italiana* | OTU35 | x | | | | | |
| **study plot 4** | | | | | | | | | | | |
| | | | VancurovaA545 | *Asterochloris phycobiontica* | OTU35 | x | | | | | |
| | | | VancurovaA546 | *Asterochloris lobophora* | OTU35 | | x | | | | |
| | | | VancurovaA547 | *Asterochloris phycobiontica* | OTU35 | | x | | | | |
| | | | VancurovaA548 | *Asterochloris phycobiontica* | OTU35 | x | | | | | |
| | | | VancurovaA549 | *Asterochloris phycobiontica* | OTU35 | | | x | | | x |
| | | | VancurovaA549V | *Asterochloris phycobiontica* | OTU35 | | | x | | | x |
| | | | VancurovaA550 | *Asterochloris phycobiontica* | OTU35 | | x | | | | |
| | | | VancurovaA552 | *Asterochloris lobophora* | OTU35 | | x | | | | |
| | | | VancurovaA553 | *Asterochloris StA5* | OTU35 | | x | | | | |
| | | | VancurovaA554V | *Asterochloris StA5* | OTU35 | | | | | | |
| | | | VancurovaA555 | *Asterochloris phycobiontica* | OTU35 | | x | | | | |
| | | | VancurovaA556 | *Asterochloris phycobiontica* | OTU35 | | | x | | | |
| **study plot 5** | | | | | | | | | | | |
| | | | VancurovaA557 | *Asterochloris StA5* | OTU35 | | x | | | | |
| | | | VancurovaA558 | *Asterochloris phycobiontica* | OTU35 | | | x | | | |
| | | | VancurovaA559 | *Asterochloris lobophora* | OTU35 | | | x | | | |
| | | | VancurovaA560 | *Asterochloris irregularis* | OTU35 | | | x | x | | |
| | | | VancurovaA561 | *Asterochloris phycobiontica* | OTU35 | x | | | | | |
| | | | VancurovaA562 | *Asterochloris irregularis* | OTU35 | | | x | | | x |
| | | | VancurovaA563V | *Asterochloris StA5* | OTU35 | | x | x | x | | |
| | | | VancurovaA564 | *Asterochloris echinata* | OTU35 | x | | | | | |
| | | | VancurovaA565 | *Asterochloris StA5* | OTU35 | | | x | | | x |
| | | | VancurovaA566 | *Asterochloris StA4* | OTU35 | x | | | | | |
| | | | VancurovaA567V | *Asterochloris phycobiontica* | OTU35 | x | | x | | | |
| | | | VancurovaA568V | *Asterochloris irregularis* | OTU35 | x | | | | | |
| | | | VancurovaA569 | *Asterochloris irregularis* | OTU35 | | | x | | | |
| **study plot 6** | | | | | | | | | | | |
| | | | VancurovaA571 | *Asterochloris StA5* | OTU35 | | | x | | | |
| | | | VancurovaA572 | *Asterochloris aff. italiana* | OTU35 | | | | x | | |
| | | | VancurovaA573 | *Asterochloris aff. italiana* | OTU35 | | | | x | | |
| | | | VancurovaA574 | *Coccomyxa viridis* | OTU35 | x | | | | | |
| | | | VancurovaA574.1 | *Elliptochloris sp.* | OTU35 | x | | | | | |
| | | | VancurovaA575 | *Asterochloris StA5* | OTU35 | | | | x | | x |
| | | | VancurovaA575.1 | *Asterochloris StA5* | OTU35 | | | | x | | x |

**Table S4** cont.

| algal ITS accession | fungal ITS accession | actin accession | sample ID | phycobiont | mycobiont | substrate/surroundings | | | | | |
|---|---|---|---|---|---|---|---|---|---|---|---|
| | | | | | | gravel | soil | moss | stone | needles | tree/shrub plants |
| | | | VancurovaA575.2 | *Asterochloris StA5* | OTU35 | | | | x | | x |
| | | | VancurovaA576 | *Asterochloris irregularis* | OTU35 | x | | | | | |
| | | | VancurovaA577 | *Asterochloris StA5* | OTU35 | x | | x | | | x |
| | | | VancurovaA578 | *Asterochloris StA5* | OTU35 | | | x | | | |
| | | | VancurovaA579 | *Asterochloris StA5* | OTU35 | | | x | | | |
| | | | VancurovaA580 | *Asterochloris StA5* | OTU35 | | | | x | | |
| | | | VancurovaA581 | *Asterochloris StA5* | OTU35 | | | | x | | |
| **study plot 7** | | | | | | | | | | | |
| | | | VancurovaA582 | *Asterochloris lobophora* | OTU35 | | | x | | | |
| | | | VancurovaA583 | *Asterochloris StA5* | OTU35 | | | x | | | |
| | | | VancurovaA584 | *Asterochloris StA5* | OTU35 | | | | x | | |
| | | | VancurovaA585 | *Asterochloris StA5* | OTU35 | | | x | | | |
| | | | VancurovaA586 | *Asterochloris clade12* | OTU35 | | | | x | | |
| | | | VancurovaA587 | *Asterochloris clade8* | OTU35 | | | x | | | |
| | | | VancurovaA588 | *Asterochloris StA5* | OTU35 | | | x | | | |
| | | | VancurovaA589 | *Asterochloris StA5* | OTU35 | | | x | | | |
| | | | VancurovaA590 | *Asterochloris StA5* | OTU35 | | | x | | | |
| | | | VancurovaA591 | *Asterochloris clade8* | OTU35 | | x | | | | |
| **study plot 8** | | | | | | | | | | | |
| | | | VancurovaA633 | *Trebouxiophyceae URa28* | OTU2 | | x | | | | |
| | | | VancurovaA634 | *Trebouxiophyceae URa28* | OTU2 | | x | x | | | |
| | | | VancurovaA634V | *Trebouxiophyceae URa28* | OTU2 | | x | x | | | |
| | | | VancurovaA635 | *Asterochloris clade8* | OTU35 | | x | | | | |
| | | | VancurovaA636 | *Asterochloris clade8* | OTU35 | | | x | | | |
| | | | VancurovaA637 | *Asterochloris phycobiontica* | OTU35 | | x | | | | |
| | | | VancurovaA638 | *Asterochloris StA5* | OTU35 | | | x | | | |
| | | | VancurovaA639 | *Asterochloris StA5* | OTU35 | | x | | | | |
| | | | VancurovaA640 | *Asterochloris phycobiontica* | OTU35 | | | x | | | |
| | | | VancurovaA641 | *Asterochloris StA5* | OTU35 | | | x | | | |
| | | | VancurovaA642 | *Asterochloris StA5* | OTU35 | | | x | | | |
| | | | VancurovaA643 | *Asterochloris phycobiontica* | OTU35 | | | x | | | |
| | | | VancurovaA644 | *Asterochloris phycobiontica* | OTU35 | | | x | | | |
| **study plot 9** | | | | | | | | | | | |
| | | | VancurovaA623 | *Asterochloris phycobiontica* | OTU35 | | | x | | | |
| | | | VancurovaA624 | *Asterochloris StA5* | OTU35 | | | x | | | |
| | | | VancurovaA625 | *Asterochloris StA5* | OTU35 | | | x | | | |
| | | | VancurovaA626 | *Asterochloris phycobiontica* | OTU35 | | | x | | | |

**Table S4** cont.

| algal ITS accession | fungal ITS accession | actin accession | sample ID | phycobiont | mycobiont | substrate/surroundings gravel | soil | moss | stone | needles | tree/shrub | plants |
|---|---|---|---|---|---|---|---|---|---|---|---|---|
| | | | VancurovaA627 | *Asterochloris StA5* | OTU35 | | | x | | | | |
| | | | VancurovaA628 | *Asterochloris StA5* | OTU35 | | | x | | | | |
| | | | VancurovaA629 | *Asterochloris lobophora* | OTU35 | | | x | | | | |
| | | | VancurovaA630 | *Asterochloris phycobiontica* | OTU35 | | | x | | | | |
| | | | VancurovaA631 | *Asterochloris phycobiontica* | OTU35 | | | x | | | | |
| | | | VancurovaA632 | *Asterochloris irregularis* | OTU35 | | | x | | | | |
| **study plot 10** | | | | | | | | | | | | |
| | | | VancurovaA645 | *Asterochloris StA5* | OTU35 | | | | x | | | |
| | | | VancurovaA646 | *Asterochloris leprarii* | OTU35 | | x | | x | | | |
| | | | VancurovaA647 | *Asterochloris StA5* | OTU35 | | x | | | | | x |
| | | | VancurovaA648 | *Asterochloris phycobiontica* | OTU35 | | | | x | | | |
| | | | VancurovaA649 | *Asterochloris StA5* | OTU35 | | | x | | | | |
| | | | VancurovaA650 | *Asterochloris phycobiontica* | OTU35 | | | x | | | | |
| | | | VancurovaA651 | *Asterochloris phycobiontica* | OTU35 | | x | | | | | |
| | | | VancurovaA652 | *Asterochloris phycobiontica* | OTU35 | | x | x | | | | |
| | | | VancurovaA653 | *Asterochloris phycobiontica* | OTU35 | | | x | x | | | |
| | | | VancurovaA654V | *Asterochloris StA5* | OTU35 | x | x | | | | | |
| | | | VancurovaA655 | *Asterochloris phycobiontica* | OTU35 | | x | | x | | | |
| **study plot 11** | | | | | | | | | | | | |
| | | | VancurovaA592 | *Asterochloris phycobiontica* | OTU35 | | | x | | | | |
| | | | VancurovaA593 | *Asterochloris StA5* | OTU35 | | | x | | | | |
| | | | VancurovaA594 | *Asterochloris StA5* | OTU35 | | | x | | | | |
| | | | VancurovaA595 | *Asterochloris StA5* | OTU35 | | | x | | | | |
| | | | VancurovaA596 | *Asterochloris StA5* | OTU35 | | | x | | | | |
| | | | VancurovaA600 | *Asterochloris italiana* | OTU2 | | | x | | | | x |
| **study plot 12** | | | | | | | | | | | | |
| | | | VancurovaA601 | *Asterochloris phycobiontica* | OTU35 | | | | x | | | |
| | | | VancurovaA602 | *Asterochloris clade8* | OTU35 | x | | | | | | |
| | | | VancurovaA603 | *Asterochloris StA4* | OTU35 | | x | | | | | |
| | | | VancurovaA604 | *Asterochloris phycobiontica* | unknown | | | | x | | | |
| | | | VancurovaA605 | *Asterochloris phycobiontica* | OTU35 | x | | | | | | |
| | | | VancurovaA606 | *Asterochloris clade12* | OTU35 | | | x | | | | |
| | | | VancurovaA607 | *Asterochloris StA5* | OTU35 | | | x | | | | |
| | | | VancurovaA608 | *Asterochloris StA5* | OTU35 | | x | | | | x | |
| | | | VancurovaA609 | *Asterochloris StA5* | OTU35 | | | x | | | | x |
| | | | VancurovaA610 | *Asterochloris phycobiontica* | OTU35 | | | | | | | x |
| | | | VancurovaA611 | *Asterochloris phycobiontica* | OTU35 | | | | x | | x | |

**Table S4** cont.

| algal ITS accession | fungal ITS accession | actin accession | sample ID | phycobiont | mycobiont | substrate/surroundings | | | | | | |
|---|---|---|---|---|---|---|---|---|---|---|---|---|
| | | | | | | gravel | soil | moss | stone | needles | tree/shrub | plants |
| **study plot 13** | | | | | | | | | | | | |
| | | | VancurovaA612 | *Asterochloris StA3* | OTU35 | | | x | | | | x |
| | | | VancurovaA613 | *Asterochloris phycobiontica* | unknown | | x | | | x | | |
| | | | VancurovaA614 | *Asterochloris StA3* | OTU35 | | x | | | x | | |
| | | | VancurovaA615 | *Asterochloris StA3* | OTU35 | | | | | | x | x |
| | | | VancurovaA616 | *Asterochloris phycobiontica* | OTU35 | | | x | | | | |
| | | | VancurovaA617 | *Asterochloris clade8* | OTU35 | | | | | x | x | |
| | | | VancurovaA618 | *Asterochloris StA3* | OTU35 | | | | | | | x |
| | | | VancurovaA619 | *Asterochloris phycobiontica* | OTU35 | | x | | | | x | |
| | | | VancurovaA620 | *Asterochloris StA3* | OTU35 | | x | | | x | | |
| | | | VancurovaA621 | *Asterochloris phycobiontica* | OTU35 | | | x | | | | |
| | | | VancurovaA622 | *Asterochloris StA5* | OTU35 | | | | x | | | |

**Table S5** Substitution models selected for each partition of *Asterochloris* and *Stereocaulon* (mycobiont) datasets and for algal ITS2 dataset using the Bayesian information criterion (BIC) as implemented in JModelTest2 (Guindon & Gascuel 2003, Darriba et al. 2012)

| **partition** | ***Asterochloris* dataset** | ***Stereocaulon* mycobiont dataset** | **Illumina metabarcoding dataset** |
|---|---|---|---|
| ITS1 rDNA | K2 + Γ (α=0.52) | T92 + Γ (α=0.6) | |
| 5.8 S rDNA | JC | JC | |
| ITS2 rDNA | K2 + Γ (α=0.27) | T92 + Γ (α=0.65) | K2 + Γ (α=0.47) |
| intron 206 of actin type I gene | T92 + Γ (α=2.49) | | |
| exon part of actin type I gene | K2 + Γ (α=0.22) | | |
| intron 248 of actin type I gene | JC | | |

**Table S6** Accession numbers of *Asterochloris* reference sequences retrieved from GenBank

| ITS accession | actin accession | sample ID | species-level lineage |
|---|---|---|---|
| AM905998 | AM906026 | Peksa 498 | *Asterochloris glomerata* |
| MH415427 | MH382148 | VancurovaO75 | *Asterochloris glomerata* |
| AF345382 | AM906024 | UTEX 895 | *Asterochloris glomerata* |
| MH415370 | MH382143 | VancurovaL992 | *Asterochloris irregularis* |
| AF345411 | AM906027 | UTEX 2236 | *Asterochloris irregularis* |
| MH415367 | MH382142 | VancurovaL988 | *Asterochloris aff. irregularis* |
| DQ229885 | DQ229888 | Talbot 281 | *Asterochloris aff. irregularis* |
| AM906012 | AM906041 | UTEX902 | *Asterochloris magna* |
| AF345440 | AM906018 | UTEX911 | *Asterochloris erici* |
| MH415428 | MH382149 | VancurovaO76 | StA1 |
| HE803036 | MH382118 | IH23 | I2 |
| MH415296 | MH382135 | VancurovaA504 | |
| HE803038 | KP318682 | IH20 | clade 9 |
| DQ229884 | DQ229896 | Nelsen 2181b | S1 |
| EU008684 | EU008711 | L54 | |
| FM945358 | FM955674 | Peksa 796 | clade 8 |
| FM945380 | FM955675 | Peksa 787 | clade 8 |
| MH415257 | MH382129 | VancurovaA378 | StA2 |
| MH415269 | MH382131 | VancurovaA392 | StA2 |
| DQ229886 | DQ229897 | Talbot KIS 187 | S3 |
| AM900492 | AM906045 | Bayerová 3401 | *Asterochloris woessiae* |
| MH415238 | MH382127 | VancurovaA337 | *Asterochloris woessiae* |
| MH415438 | MH382150 | VancurovaO98 | StA3 |
| KX051235 | KX051239 | KGS007A | *Asterochloris sejongensis* |
| KX051236 | KX051240 | KGS064B | *Asterochloris sejongensis* |
| MH415374 | MH382144 | VancurovaO10 | clade 12 |
| FM945378 | FM955677 | Peksa 921 | clade 12 |
| DQ229877 | DQ229898 | Nelsen 3974 | *Asterochloris friedlii* |
| AM905995 | AM906021 | Peksa 235 | *Asterochloris friedlii* |
| HE803033 | MH382119 | IH31 | I1 |
| HE803029 | MH382117 | I6 | I1 |
| KP257384 | KP257351 | C19 | *Asterochloris mediterranea* |
| EU008690 | EU008715 | L60 | URa14 |
| MH415220 | MH382124 | VancurovaA14 | |
| MH415422 | MH382147 | VancurovaO70 | *Asterochloris aff. italiana* |
| MH415217 | MH382121 | VancurovaA10 | *Asterochloris italiana* |
| AM906001 | AM906030 | CCAP 219/5B | *Asterochloris italiana* |
| MH415329 | MH382137 | VancurovaL1074 | StA4 |
| DQ229882 | DQ229893 | Talbot 400 | StA4 |
| AM900490 | AM906042 | SAG 26.81 | *Asterochloris phycobiontica* |
| MH415414 | MH382146 | VancurovaO50 | StA5 |
| MH415366 | MH382141 | VancurovaL958 | StA5 |
| FN556044 | KP318679 | Peksa 866 | *Asterochloris lobophora* |
| MH415307 | MH382136 | VancurovaDS3.1 | *Asterochloris lobophora* |
| AM906008 | AM906037 | Peksa 166 | *Asterochloris lobophora* |
| KP318676 | KP318681 | Peksa 495 | A4 |
| AM905992 | AM906017 | Peksa 186 | *Asterochloris echinata* |
| FM955667 | FM955671 | Peksa 551 | *Asterochloris echinata* |
| AM905993 | AM906019 | UTEX 1714 | *Asterochloris excentrica* |
| AM906002 | AM906031 | Peksa 183 | *Asterochloris leprarii* |
| FN556035 | FN556048 | Peksa 860 | A9 |
| MH415218 | MH382122 | VancurovaA11 | A9 |
| MH415216 | MH382120 | VancurovaA1 | StA6 |
| MH415219 | MH382123 | VancurovaA13 | StA7 |
| MH415287 | MH382132 | VancurovaA496 | StA8 |
| MH415288 | MH382133 | VancurovaA498 | StA8 |
| FM955669 | FM955673 | Peksa 900 | *Asterochloris gaertneri* |
| AM905997 | AM906023 | Peksa 236 | *Asterochloris gaertneri* |
| FN556042 | FN556051 | Peksa 873 | A11 |
| FN556043 | FN556052 | Peksa 870 | A11 |

**Table S7** Relative abundance of each ASV in each *Stereocaulon* sample

| ASV | A523M | A554M | A563M | A570M | A597M | A598M | A634M | A633M |
|---|---|---|---|---|---|---|---|---|
| ***Asterochloris*** | **96,41%** | **91,50%** | **98,77%** | **52,44%** | **0,00%** | **9,90%** | **6,19%** | **7,41%** |
| 0773d605c4043cdd107243001c6490e6 | 0,39% | 0,00% | 0,00% | 0,00% | 0,00% | 0,00% | 0,00% | 0,00% |
| 22feecfeebfed298019ccc2de7ea7a4f | 82,80% | 0,00% | 0,00% | 0,51% | 0,00% | 0,00% | 0,00% | 0,00% |
| 4e1d7c47e2c4c3b51fd0746d0bdc4d77 | 0,00% | 0,00% | 0,00% | 0,00% | 0,00% | 0,00% | 0,00% | 0,17% |
| 7def077ab2105529e2da667181b45321 | 0,00% | 0,50% | 0,00% | 0,00% | 0,00% | 0,00% | 0,00% | 0,00% |
| 85be31a9a9656c804dfaa8d63f0b4c86 | 0,00% | 0,38% | 0,00% | 0,00% | 0,00% | 0,00% | 1,01% | 4,18% |
| 87ac6b49cc5d1ee84b1b41184fe3ac17 | 0,00% | 0,00% | 0,00% | 0,24% | 0,00% | 0,00% | 0,00% | 0,00% |
| a49f5abdcab6d47686f7384090d0b98f | 0,00% | 0,00% | 0,00% | 0,21% | 0,00% | 0,00% | 0,00% | 0,00% |
| a4e586faa69d099f647fbbf7c5c781d8 | 0,88% | 90,62% | 10,08% | 50,42% | 0,00% | 0,00% | 0,37% | 0,00% |
| aad656bc9e55af9ff1f14fda4c7331d4 | 0,00% | 0,00% | 0,00% | 0,48% | 0,00% | 0,00% | 0,00% | 0,00% |
| d48be1abd9465c864d8e54ee080c1864 | 2,65% | 0,00% | 88,69% | 0,58% | 0,00% | 4,14% | 0,66% | 0,00% |
| dfd67be0a040eddaa59f02bab8503d01 | 0,00% | 0,00% | 0,00% | 0,00% | 0,00% | 0,00% | 0,00% | 0,48% |
| e99727d63dd17ec15ae526ed06e91141 | 9,69% | 0,00% | 0,00% | 0,00% | 0,00% | 0,00% | 0,00% | 0,00% |
| fbbb323f92a953f33f0d1c2bbd751388 | 0,00% | 0,00% | 0,00% | 0,00% | 0,00% | 0,00% | 4,15% | 2,59% |
| 1090b5b3f02a9f90e97674d9db6c8a21 | 0,00% | 0,00% | 0,00% | 0,00% | 0,00% | 0,28% | 0,00% | 0,00% |
| 36bce30657ed4c3a02ad3c94ce7189cf | 0,00% | 0,00% | 0,00% | 0,00% | 0,00% | 1,01% | 0,00% | 0,00% |
| 7c06c79b6591c393dd587c064bb272ac | 0,00% | 0,00% | 0,00% | 0,00% | 0,00% | 3,32% | 0,00% | 0,00% |
| 873373e1b12419297c358f3d3f05f3c6 | 0,00% | 0,00% | 0,00% | 0,00% | 0,00% | 1,15% | 0,00% | 0,00% |
| **Clade VII** | **0,00%** | **0,00%** | **0,00%** | **0,00%** | **0,00%** | **0,00%** | **0,00%** | **0,76%** |
| 6640f588c3c03e787f910baed715cd75 | 0,00% | 0,00% | 0,00% | 0,00% | 0,00% | 0,00% | 0,00% | 0,76% |
| **Clade XVIII** | **0,00%** | **0,00%** | **0,00%** | **45,01%** | **0,75%** | **9,33%** | **0,00%** | **0,00%** |
| 22f0d7ffbb54a41fbaed575bbf236074 | 0,00% | 0,00% | 0,00% | 2,03% | 0,14% | 0,83% | 0,00% | 0,00% |
| 33277e83aaf07d90bf2b7df2f8c6c526 | 0,00% | 0,00% | 0,00% | 14,57% | 0,00% | 0,00% | 0,00% | 0,00% |
| 363eb19dabdce2db62981f298e58d38a | 0,00% | 0,00% | 0,00% | 1,27% | 0,00% | 0,00% | 0,00% | 0,00% |
| 5a78297855b6a01b7025236c3ed33970 | 0,00% | 0,00% | 0,00% | 0,36% | 0,00% | 0,00% | 0,00% | 0,00% |
| 693e873a1e85633a5e847c2813b848f5 | 0,00% | 0,00% | 0,00% | 10,36% | 0,00% | 0,00% | 0,00% | 0,00% |
| 7f5e0f713269e9b0e05b68775e45a748 | 0,00% | 0,00% | 0,00% | 0,31% | 0,00% | 0,00% | 0,00% | 0,00% |
| 8297ec6dfe58d45468b7e8e47dffcbba | 0,00% | 0,00% | 0,00% | 0,00% | 0,49% | 4,68% | 0,00% | 0,00% |
| 93feaa8c6fc846943a590cdf36fcc6bd | 0,00% | 0,00% | 0,00% | 2,73% | 0,00% | 0,00% | 0,00% | 0,00% |
| a34fcf4ef83bd7a07368b48914cb0bf6 | 0,00% | 0,00% | 0,00% | 1,19% | 0,00% | 0,00% | 0,00% | 0,00% |
| af66fe2037bb72fa617f606e31547fa4 | 0,00% | 0,00% | 0,00% | 0,28% | 0,00% | 0,00% | 0,00% | 0,00% |
| cbd6b0959a3b740b29299a82d9c7fd54 | 0,00% | 0,00% | 0,00% | 8,86% | 0,00% | 0,00% | 0,00% | 0,00% |
| d743cedc743576ac50e4f0f6333a26cc | 0,00% | 0,00% | 0,00% | 0,00% | 0,11% | 0,00% | 0,00% | 0,00% |
| d776d518d8060add1e1245191d831d4f | 0,00% | 0,00% | 0,00% | 3,05% | 0,00% | 1,44% | 0,00% | 0,00% |
| fa5385b7d826cf1ef315a3bb60ae6204 | 0,00% | 0,00% | 0,00% | 0,00% | 0,00% | 1,71% | 0,00% | 0,00% |
| ee3931af80112ff916a60510bde76faf | 0,00% | 0,00% | 0,00% | 0,00% | 0,00% | 0,67% | 0,00% | 0,00% |
| ***Coccomyxa*** | **0,00%** | **0,00%** | **0,00%** | **0,19%** | **0,24%** | **0,00%** | **0,77%** | **1,14%** |
| 1a98adae19579aaf9ff1dc9966dd90d7 | 0,00% | 0,00% | 0,00% | 0,00% | 0,24% | 0,00% | 0,00% | 0,00% |
| 66bdbfa7f8cbe36a07a9ec130b8c5beb | 0,00% | 0,00% | 0,00% | 0,19% | 0,00% | 0,00% | 0,18% | 0,26% |
| 91f5b6e0fae94b0ac9021eb79f36f7b6 | 0,00% | 0,00% | 0,00% | 0,00% | 0,00% | 0,00% | 0,11% | 0,00% |
| a0b8a5d4749d9250968aee293413dd3a | 0,00% | 0,00% | 0,00% | 0,00% | 0,00% | 0,00% | 0,24% | 0,14% |
| f054020cbf736e42584284620a6a2030 | 0,00% | 0,00% | 0,00% | 0,00% | 0,00% | 0,00% | 0,00% | 0,29% |
| f1f932ec5671a64311f14f51c589ea40 | 0,00% | 0,00% | 0,00% | 0,00% | 0,00% | 0,00% | 0,09% | 0,45% |
| fd2da8fccdad584a8d294151d71376f9 | 0,00% | 0,00% | 0,00% | 0,00% | 0,00% | 0,00% | 0,15% | 0,00% |
| ***Coelastrella*** | **0,00%** | **0,00%** | **0,00%** | **0,00%** | **0,00%** | **0,00%** | **0,31%** | **2,17%** |
| cd98ddfaddf0c563b533df10b83c94f2 | 0,00% | 0,00% | 0,00% | 0,00% | 0,00% | 0,00% | 0,00% | 1,67% |
| edf68c3189fe6efa4e4a7ce1b18fcd54 | 0,00% | 0,00% | 0,00% | 0,00% | 0,00% | 0,00% | 0,13% | 0,00% |
| fae8d63a97b2686ab9b1683a82f34a24 | 0,00% | 0,00% | 0,00% | 0,00% | 0,00% | 0,00% | 0,18% | 0,50% |
| ***Coenochloris*** | **0,00%** | **0,25%** | **0,00%** | **0,00%** | **0,00%** | **0,00%** | **0,00%** | **0,00%** |
| 36490d68e1cc6ab6ad5b518f09870a97 | 0,00% | 0,25% | 0,00% | 0,00% | 0,00% | 0,00% | 0,00% | 0,00% |
| ***Elliptochloris*** | **0,00%** | **0,41%** | **0,00%** | **0,43%** | **0,26%** | **0,00%** | **2,22%** | **6,51%** |
| 033517ce8a44175727c4b60b9243a922 | 0,00% | 0,00% | 0,00% | 0,00% | 0,00% | 0,00% | 0,00% | 0,19% |
| 055a30c59b0804cb5ed7827d5ea0d5d1 | 0,00% | 0,00% | 0,00% | 0,00% | 0,00% | 0,00% | 0,17% | 0,18% |
| 15af57f2867287f7cc8e437f1a48660d | 0,00% | 0,00% | 0,00% | 0,00% | 0,00% | 0,00% | 0,00% | 0,26% |
| 277091179c651030797c72b0f84539d8 | 0,00% | 0,00% | 0,00% | 0,14% | 0,13% | 0,00% | 0,00% | 0,00% |
| 29caf6696a5a44173052920c5849786d | 0,00% | 0,00% | 0,00% | 0,00% | 0,00% | 0,00% | 0,07% | 0,00% |
| 2a7edb47da012d25dc9fe74f0895cf3f | 0,00% | 0,00% | 0,00% | 0,00% | 0,00% | 0,00% | 0,00% | 0,26% |
| 2d288fda32d8a03001f88926b1c9d84f | 0,00% | 0,00% | 0,00% | 0,29% | 0,00% | 0,00% | 0,00% | 0,00% |

**Table S7** cont.

| ASV | A523M | A554M | A563M | A570M | A597M | A598M | A634M | A633M |
|---|---|---|---|---|---|---|---|---|
| 34238f9f2d09f5306a7cabcf8e4b9ce5 | 0,00% | 0,00% | 0,00% | 0,00% | 0,00% | 0,00% | 0,11% | 0,00% |
| 37c413557e64a9064c3fc68c11a7421a | 0,00% | 0,00% | 0,00% | 0,00% | 0,00% | 0,00% | 0,15% | 0,00% |
| 5081b1190bb69e7adbb9a58169c647e9 | 0,00% | 0,00% | 0,00% | 0,00% | 0,00% | 0,00% | 0,00% | 0,26% |
| 59e46a1330290f14f1f13c981cf5b13a | 0,00% | 0,00% | 0,00% | 0,00% | 0,00% | 0,00% | 0,00% | 0,31% |
| 5aa3a8de6e711d0706d17fc32712d3dd | 0,00% | 0,00% | 0,00% | 0,00% | 0,00% | 0,00% | 0,70% | 0,73% |
| 5fd4222aff69b399a4df046a41d816e3 | 0,00% | 0,00% | 0,00% | 0,00% | 0,00% | 0,00% | 0,00% | 0,78% |
| 6dd16d34f54043bf18bbae77afb6cc2d | 0,00% | 0,00% | 0,00% | 0,00% | 0,00% | 0,00% | 0,00% | 0,71% |
| 858042d7fc87331b567dab4162fbd311 | 0,00% | 0,00% | 0,00% | 0,00% | 0,13% | 0,00% | 0,00% | 0,00% |
| 95c03e83804c9613c5b8afbe71965c26 | 0,00% | 0,41% | 0,00% | 0,00% | 0,00% | 0,00% | 0,44% | 0,15% |
| a33566bfe4b0d2b16e2f4523874da2fd | 0,00% | 0,00% | 0,00% | 0,00% | 0,00% | 0,00% | 0,00% | 0,52% |
| a667fa3f885abd611cf0f297f60c0355 | 0,00% | 0,00% | 0,00% | 0,00% | 0,00% | 0,00% | 0,00% | 0,21% |
| a9b4e6f9ed5388afa56a256426f3cd7e | 0,00% | 0,00% | 0,00% | 0,00% | 0,00% | 0,00% | 0,00% | 0,24% |
| bfe815229d41f5376cd18ef7f38b88cb | 0,00% | 0,00% | 0,00% | 0,00% | 0,00% | 0,00% | 0,00% | 0,15% |
| e1f903aad02e5332261a5c28eacebb0f | 0,00% | 0,00% | 0,00% | 0,00% | 0,00% | 0,00% | 0,00% | 0,21% |
| f7571040bdf6f5cba93fe4ec831f30d6 | 0,00% | 0,00% | 0,00% | 0,00% | 0,00% | 0,00% | 0,13% | 0,00% |
| fc8d038ce995a2d3389256592c1c201d | 0,00% | 0,00% | 0,00% | 0,00% | 0,00% | 0,00% | 0,46% | 1,35% |
| ***Chloroidium*** | **0,00%** | **0,00%** | **0,00%** | **0,20%** | **0,07%** | **0,65%** | **0,53%** | **1,05%** |
| 12a64454e2a9d725208dec06fa167461 | 0,00% | 0,00% | 0,00% | 0,20% | 0,00% | 0,00% | 0,00% | 0,00% |
| 23666d2107e2329e3f34ad474919df44 | 0,00% | 0,00% | 0,00% | 0,00% | 0,00% | 0,65% | 0,00% | 0,00% |
| 761496fb7ace9bdab1accbab50ff917c | 0,00% | 0,00% | 0,00% | 0,00% | 0,00% | 0,00% | 0,53% | 0,00% |
| d5ec63a3b55a952369c5c8254d0132df | 0,00% | 0,00% | 0,00% | 0,00% | 0,07% | 0,00% | 0,00% | 1,05% |
| ***Lobosphaera*** | **0,00%** | **0,00%** | **0,00%** | **0,00%** | **0,00%** | **0,00%** | **0,08%** | **2,45%** |
| 247289abba6262468dec4281db6ef9e7 | 0,00% | 0,00% | 0,00% | 0,00% | 0,00% | 0,00% | 0,08% | 0,00% |
| 419cf5613cac948aceace26cecd90c5c | 0,00% | 0,00% | 0,00% | 0,00% | 0,00% | 0,00% | 0,00% | 0,74% |
| 8926000fbcb4214c4f3a178884e0c857 | 0,00% | 0,00% | 0,00% | 0,00% | 0,00% | 0,00% | 0,00% | 0,31% |
| 90a5cde57da67ec7fa8ea6ad33f6e771 | 0,00% | 0,00% | 0,00% | 0,00% | 0,00% | 0,00% | 0,00% | 0,37% |
| c435a0466fb6fa01287354c8a1dfe783 | 0,00% | 0,00% | 0,00% | 0,00% | 0,00% | 0,00% | 0,00% | 0,45% |
| fe80971a622f0242474809f6ab26b01d | 0,00% | 0,00% | 0,00% | 0,00% | 0,00% | 0,00% | 0,00% | 0,57% |
| ***Myrmecia*** | **0,00%** | **0,00%** | **0,00%** | **0,00%** | **0,00%** | **0,00%** | **0,09%** | **0,97%** |
| 58e7be6488435b4d3eb92b4ded51dfb3 | 0,00% | 0,00% | 0,00% | 0,00% | 0,00% | 0,00% | 0,09% | 0,97% |
| ***Neocystis*** | **0,00%** | **0,00%** | **0,00%** | **0,00%** | **0,00%** | **0,00%** | **0,00%** | **0,59%** |
| 30d80e23cdbd5dd56244611e687a83ab | 0,00% | 0,00% | 0,00% | 0,00% | 0,00% | 0,00% | 0,00% | 0,25% |
| 5999dd175f2cbeed671a906d82ab4071 | 0,00% | 0,00% | 0,00% | 0,00% | 0,00% | 0,00% | 0,00% | 0,20% |
| 6a13d1aa391c13110b7872403bd38ad1 | 0,00% | 0,00% | 0,00% | 0,00% | 0,00% | 0,00% | 0,00% | 0,13% |
| ***Pseudochlorella*** | **0,40%** | **0,31%** | **0,00%** | **0,00%** | **2,59%** | **1,25%** | **0,29%** | **4,15%** |
| 231db3d3289f45f0258fb36841e7a5f3 | 0,00% | 0,00% | 0,00% | 0,00% | 1,56% | 0,00% | 0,00% | 0,00% |
| 26807efe882be5c05b8ba4cf090160d9 | 0,00% | 0,00% | 0,00% | 0,00% | 0,37% | 0,55% | 0,11% | 0,40% |
| 31d06c91e033c9f93a92f6d30b82a7f2 | 0,00% | 0,00% | 0,00% | 0,00% | 0,23% | 0,35% | 0,00% | 0,00% |
| 4f58a48e04c4e0811fdd652878e78aaa | 0,00% | 0,13% | 0,00% | 0,00% | 0,29% | 0,00% | 0,00% | 2,91% |
| 521c77e3ae9b248f635599264c6a24a0 | 0,40% | 0,00% | 0,00% | 0,00% | 0,14% | 0,36% | 0,00% | 0,21% |
| 544ba515f0403962df649ee1f2d78c8a | 0,00% | 0,00% | 0,00% | 0,00% | 0,00% | 0,00% | 0,09% | 0,62% |
| d6694b81dd44c3535340a26e421c18f0 | 0,00% | 0,18% | 0,00% | 0,00% | 0,00% | 0,00% | 0,09% | 0,00% |
| ***Sanguina*** | **0,00%** | **0,00%** | **0,00%** | **1,74%** | **16,58%** | **7,80%** | **0,00%** | **0,00%** |
| 2ae5b7067248eef6014856f10392c9bb | 0,00% | 0,00% | 0,00% | 0,00% | 0,11% | 0,00% | 0,00% | 0,00% |
| 8f24e8096c8044822a68c47eea5258a6 | 0,00% | 0,00% | 0,00% | 0,00% | 1,02% | 0,00% | 0,00% | 0,00% |
| c55683ee30cbfbc1c88ff66dea615a78 | 0,00% | 0,00% | 0,00% | 0,00% | 13,95% | 7,48% | 0,00% | 0,00% |
| d536470db6babef083c90919089c7ab5 | 0,00% | 0,00% | 0,00% | 1,74% | 0,00% | 0,00% | 0,00% | 0,00% |
| eb952482b2be823acc8b920a19f152ca | 0,00% | 0,00% | 0,00% | 0,00% | 1,51% | 0,00% | 0,00% | 0,00% |
| 29e6aeb89a584ffe7d533ef201a78c56 | 0,00% | 0,00% | 0,00% | 0,00% | 0,00% | 0,32% | 0,00% | 0,00% |
| ***Trebouxia*** | **0,19%** | **0,08%** | **0,00%** | **0,00%** | **6,17%** | **0,00%** | **0,00%** | **0,00%** |
| 0f66f67d62eec5a1a146f5bad02fbe77 | 0,00% | 0,00% | 0,00% | 0,00% | 5,40% | 0,00% | 0,00% | 0,00% |
| 1076bf11d017bd0a6e68bcbf2a893dfc | 0,00% | 0,00% | 0,00% | 0,00% | 0,11% | 0,00% | 0,00% | 0,00% |
| 409a64288124ed58185eae54123f2b44 | 0,10% | 0,00% | 0,00% | 0,00% | 0,00% | 0,00% | 0,00% | 0,00% |
| 563203bee5f624f406bce0fc3c0aa45f | 0,00% | 0,00% | 0,00% | 0,00% | 0,50% | 0,00% | 0,00% | 0,00% |
| 89a3241895fd23d40c08452930898069 | 0,00% | 0,08% | 0,00% | 0,00% | 0,00% | 0,00% | 0,00% | 0,00% |
| 8d146472bc7b7a0e8357bd9e6d1108e4 | 0,09% | 0,00% | 0,00% | 0,00% | 0,00% | 0,00% | 0,00% | 0,00% |
| b4af216a92d266968878af1e6116655d | 0,00% | 0,00% | 0,00% | 0,00% | 0,10% | 0,00% | 0,00% | 0,00% |
| cd49715fbe3dd681f381593b9839be96 | 0,00% | 0,00% | 0,00% | 0,00% | 0,06% | 0,00% | 0,00% | 0,00% |

**Table S7** cont.

| ASV | A523M | A554M | A563M | A570M | A597M | A598M | A634M | A633M |
|---|---|---|---|---|---|---|---|---|
| **URa25** | **0,00%** | **0,00%** | **0,00%** | **0,00%** | **0,00%** | **0,00%** | **0,00%** | **0,63%** |
| 996b61556135c0ce99be965f27b223cf | 0,00% | 0,00% | 0,00% | 0,00% | 0,00% | 0,00% | 0,00% | 0,63% |
| **URa28** | **3,00%** | **7,44%** | **1,23%** | **0,00%** | **73,35%** | **71,07%** | **89,53%** | **72,17%** |
| 0bd320046d0690c3ce85d7d54b324056 | 0,00% | 0,00% | 0,00% | 0,00% | 0,56% | 0,00% | 0,00% | 0,00% |
| 144c9e2e50d9090ce392b99024852df2 | 0,00% | 0,00% | 0,00% | 0,00% | 0,36% | 0,00% | 0,00% | 0,00% |
| 166d7ed41986a185bb48eb6930e31656 | 0,05% | 0,15% | 0,00% | 0,00% | 1,50% | 0,50% | 0,00% | 0,00% |
| 27545897ab85f79aebb3dd70925b41da | 2,79% | 3,99% | 0,45% | 0,00% | 70,06% | 70,57% | 88,58% | 68,54% |
| 3c453af8ca7d321a75429b04f8586a15 | 0,00% | 0,00% | 0,00% | 0,00% | 0,13% | 0,00% | 0,00% | 0,00% |
| 4e5d738c8700b5fe02217f0260bb78b3 | 0,08% | 0,00% | 0,00% | 0,00% | 0,00% | 0,00% | 0,00% | 0,00% |
| 650970e3bb2e4c9cfb41c3d64098c5f1 | 0,08% | 3,05% | 0,78% | 0,00% | 0,00% | 0,00% | 0,00% | 3,63% |
| 6ac8727515be1be0a25530380ffb3715 | 0,00% | 0,07% | 0,00% | 0,00% | 0,00% | 0,00% | 0,00% | 0,00% |
| 81b9faca3807e0c02a18b4ac65e869cc | 0,00% | 0,00% | 0,00% | 0,00% | 0,00% | 0,00% | 0,95% | 0,00% |
| bec1ded9a4a1d62696263b01f4478d53 | 0,00% | 0,00% | 0,00% | 0,00% | 0,09% | 0,00% | 0,00% | 0,00% |
| c9e4620bd01c1facd9b385518fcd9340 | 0,00% | 0,00% | 0,00% | 0,00% | 0,43% | 0,00% | 0,00% | 0,00% |
| e081a3ae0c72c8e72cfd0efe91ccc6ae | 0,00% | 0,00% | 0,00% | 0,00% | 0,22% | 0,00% | 0,00% | 0,00% |
| e2ef35e8238c75c61db809a741c07713 | 0,00% | 0,18% | 0,00% | 0,00% | 0,00% | 0,00% | 0,00% | 0,00% |
