## Supplementary material for "Symbiosis between river and dry lands: phycobiont dynamics on river gravel bars": Online Resource 2

**List of contents:**

Fig. S1 Page 2

Fig. S2 Page 3

Fig. S3 Page 4

Fig. S4 Page 5

Fig. S5 Page 6

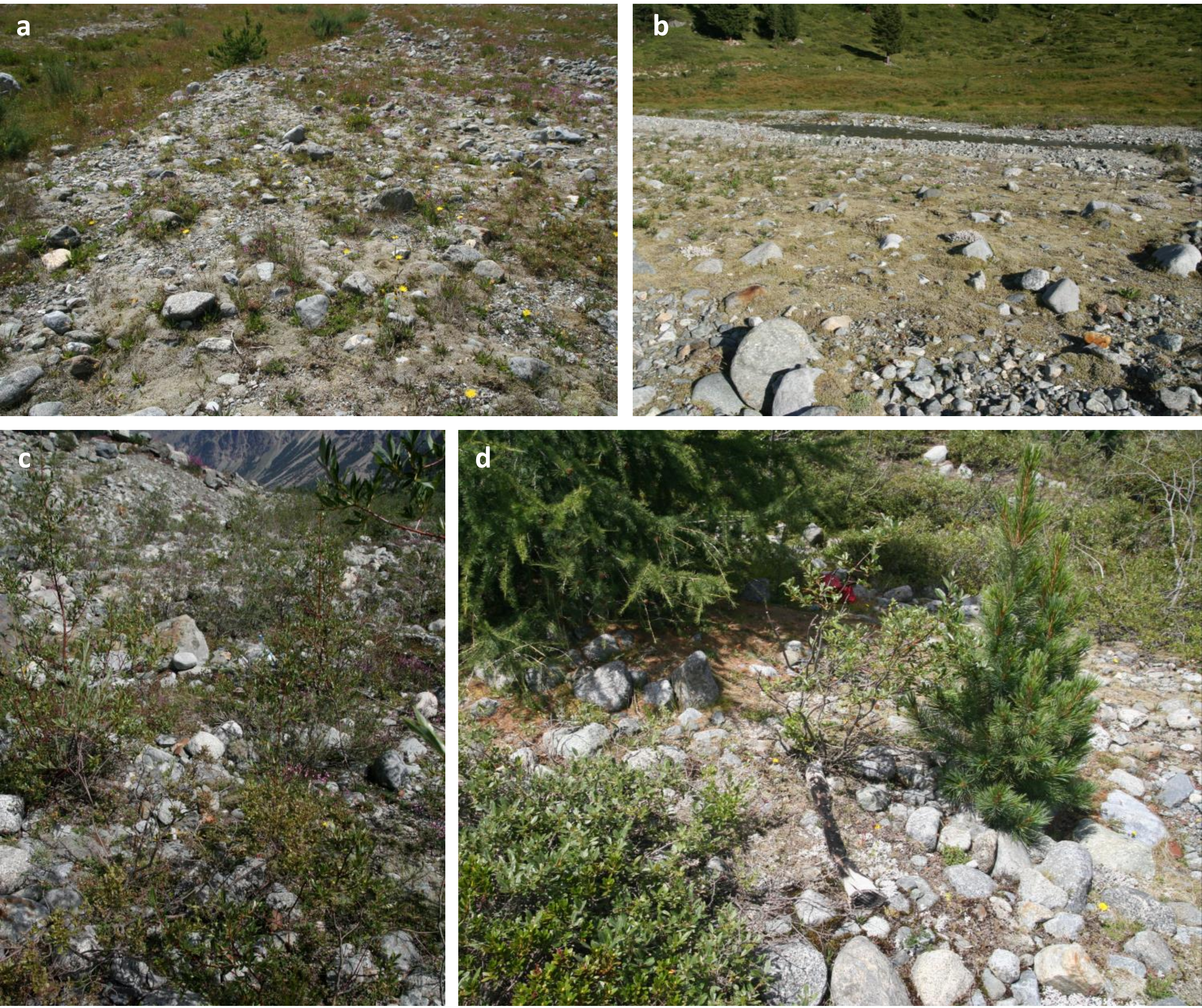


**Fig. S1** Examples of study plots belonging to different succession stages: **a, b** succession stage 1 (plots 5 and 9), **c** succession stage 2 (plot 2), **d** succession stage 3 (plot 3)

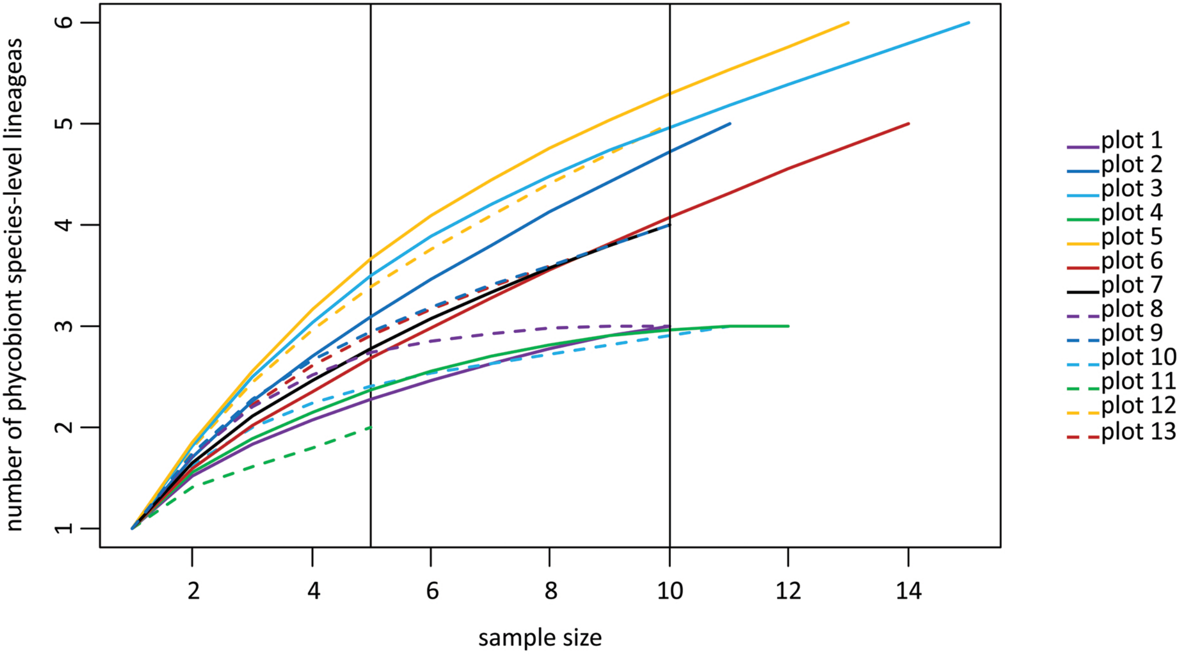


**Fig. S2** Rarefaction curves for 13 study plots. Vertical lines are drawn at sample size of n=5 and sample size of n=10

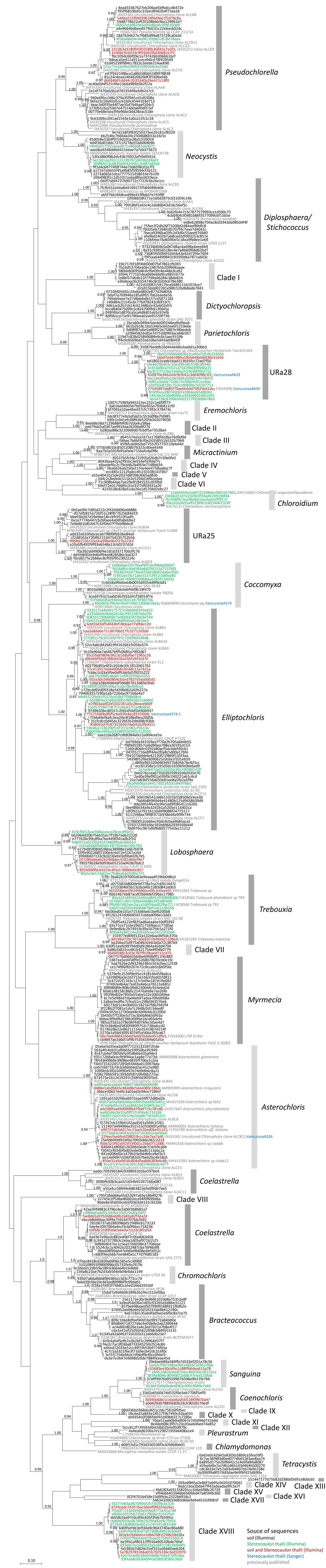


**Fig. S3** Phylogenetic hypothesis resulting from Bayesian analysis of the algal ITS2 rDNA sequences obtained by Illumina metabarcoding of soil samples, selected lichen samples and reference sequences from GenBank. Clade affiliations: URa25, URa28 *sensu* Ruprecht et al. (2014). Clades I–XVIII were identified as new in present study. Values at the nodes indicate the statistical supports of Bayesian posterior probability

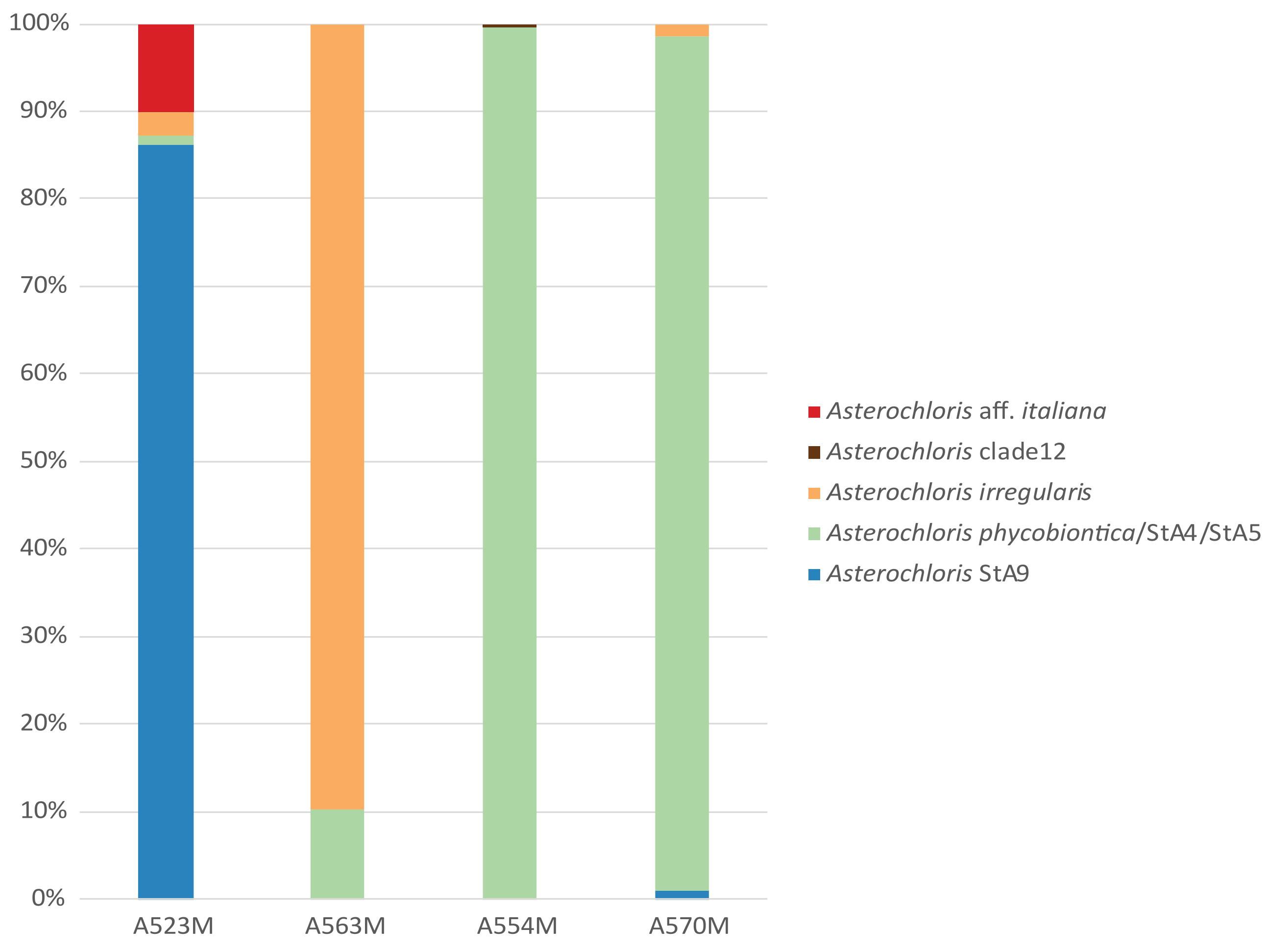


**Fig. S4** Relative frequency of ASVs linked to various *Asterochloris* species. Solely samples with *Asterochloris* as a predominant phycobiont (>50% algal reads belonged to *Asterochloris*) were displayed

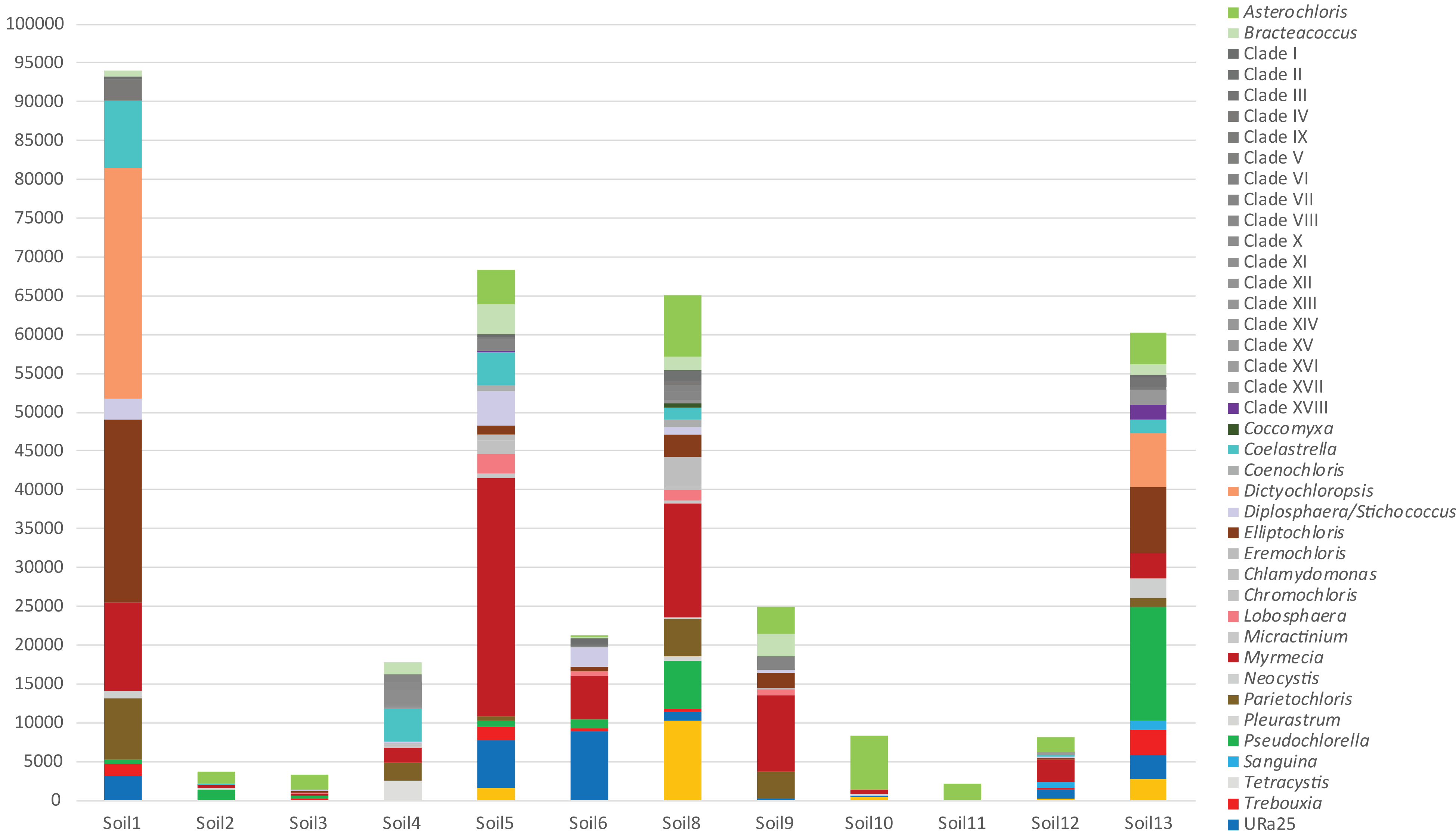


**Fig. S5** Frequency of ASVs linked to algal clades recovered from soil samples. Clade affiliations: URa25, URa28 *sensu* Ruprecht et al. (2014). Clades I–XVIII were identified as new in present study. Amplicon sequence variants (ASVs) were sorted into these clades based on phylogenetic hypothesis presented in Online Resource 2/Fig. S3
